# *C*-4-Modified Isotetrones Prevent Biofilm Growth and Persister Cell Resuscitation in *Mycobacterium smegmatis*

**DOI:** 10.1101/2022.10.23.513374

**Authors:** Kingshuk Bag, Aditya Kumar Pal, Subhadip Basu, Mamta Singla, Dipankar Chatterji, Prabal Kumar Maiti, Anirban Ghosh, Narayanaswamy Jayaraman

## Abstract

Hyperphosphorylated guanosine nucleotide (p)ppGpp, synthesized by Rel proteins, regulates the stringent response pathway responsible for biofilm growth and persister cell formation in the stationary phase of mycobacteria. The discovery of vitamin C as a potent inhibitor of Rel protein activities raises the prospect of such a tetrone lactone to prevent biofilm growth and persister cell formation. The closely related isotetrone lactone derivatives are identified in the present study as potent inhibitors of the above processes in a mycobacterium. Isotetrone lactone derivatives are synthesized from appropriate α-ketocarboxylic acids, derived from the a-amino acids. Aldol condensation with formaldehyde, followed by the lactone formation, completes synthesis of isotetrone derivatives, possessing varied substituents at *C*-4 carbon, in good yields. A series of biochemical evaluations of biofilm growth and persister cell formation in *M. smegmatis* is conducted. Among the derivatives, isotetrone possessing phenyl substituent at *C*-4 carbon completely inhibit the biofilm formation at 400 μg mL^-1^ concentration, 84 h of post-exposure, followed by a moderate inhibition by the isotetrone possessing *p*-hydroxyphenyl substituent. Whereas, the latter isotetrone inhibits the growth of cells at 400 μg mL^-1^ f.c. when monitored for 2 weeks, under PBS starvation condition. Isotetrones also potentiate the inhibition of antibiotic tolerant regrowth of cells by ciprofloxacin antibiotic (0.75 μg mL^-1^) and thus act as bio-enhancers. The combination is shown to significantly arrest the emergence of ciprofloxacin-resistant genetic mutants. The observations suggest that isotetrones in combination with ciprofloxacin are therapeutically superior when administered together. Systematic molecular dynamics studies show that isotetrone derivative binds to Rel protein more efficiently than vitamin C and the binding is aided by hydrogen bonding, van der Waals and electrostatic interactions at a binding site possessing serine, threonine, lysine and arginine residues. The present study establishes that the identified isotetrone derivatives (i) act as inhibitors of *M. smegmatis* biofilm growth and (ii) arrest the re-emergence of recalcitrant persister cells when administered together with ciprofloxacin antibiotic. Results of this study establish that isotetrones as new chemical entities that interfere with stringent response pathways in a mycobacterium under stress and permit overcoming the multidrug-resistant persister cell emergence in the bacterium.

## Introduction

Persister cells form a group of an antibiotic-tolerant bacterial cell population, responsible for the resurgence of pathogenic infections upon withdrawal of antibiotics on infected host cells.^1^ Among other factors, the stringent response pathway regulates the persister cell emergence and the pathway is regulated heavily by hyperphosphorylated guanosine nucleotides, namely, (p)ppGpp.^2–7^ Sustained investigations show the intense inter-relationship of mycobacterial stringent response pathway, persister cell growth and infection to the host cells by overcoming antibiotic treatments.^8–12^ The emergence of persister cells is often correlated to the synthesis of (p)ppGpp by Rel, which is a predominant bacterial enzyme. Under stress, such as, in starvation conditions, the Rel enzyme activates the (p)ppGpp synthesis, which, in turn, leads to persister cell growth, possessing altered metabolic processes. Targeting the stringent response pathway, and thus the (p)ppGpp synthesis, appears to be a promising approach to overcome persister cell-mediated infections to host cells.

Relacin and derivatives thereof have been developed as potent Rel inhibitors, although their *in vivo* inhibition activities occur in millimolar ranges.^13–15^ Among other potent inhibitors studied so far, vitamin C shows promising inhibitory activities by inhibition of the ppGpp synthesis.^11^ Vitamin C binds directly to the Rel enzyme and inhibits (p)ppGpp biosynthesis, thereby establishing the importance of vitamin C to prevent mycobacterial persister cell emergence. The inhibition of (p)ppGpp biosynthesis occurs at millimolar concentrations of vitamin C, although such high concentrations of vitamin C would not be an impediment to a normal course of administration to antibiotic-tolerant infected cells populated with persisters.

Vitamin C has also been discovered recently as a potent inhibitor of persister cell growth in *M*. *smegmatis* and *M*. *tuberculosis* bacterial species.^16–21^ These discoveries prompt studies of newer chemical entities possessing the tetrone lactone scaffold. Vitamin C is a tetrone lactone^22^ and thus scaffold search of potential inhibitors with a varied substituent on the tetrone lactone is a viable approach. The present report demonstrates the potencies of newly formed *C*-4 modified isotetronic acids as inhibitors of *M*. *smegmatis* persister cell growth, through a series of biochemical evaluations.

## Results and Discussion

### Chemical synthesis of C-4-Modified isotetronic acids

The a-hydroxy-g-butyrolactones, namely, isotetrone lactones form as a scaffold for many important natural products of therapeutic importance.^23–27^ Many natural products possess the a-hydroxy-g-butyrolactone and modifications around this scaffold form the structural variations that each isotetrone natural product is constituted with.^28–34^ Early biosynthesis studies by Yamamoto and coworkers demonstrated that a-amino acids form the substrate for the formation of a-hydroxy-g-butyrolactone metabolites, produced during the culture growth of fungus *Aspergillus terreus* IFO 8835 strain.^35^ The deamination of a-amino acids was presumed to form the first of several steps, leading to isotetrone formation. The deamination process led to the transformation of the a-amino acid to a-keto acid, namely, the pyruvic acid. Pyruvic acid acts as an excellent nucleophile to aldol reactions with carbonyl electrophiles and in biosynthesis, the reaction is mediated by a type II aldolase enzyme.^36–37^ A subsequent lactonization leads the resulting cross-aldol product to the isotetrone. Pyruvic acid as an important synthon is well-exploited in the chemical synthesis of many derivatives of isotetrone, with substitutions at *C*-3 to *C*-5 carbons.^38–40^Enantioselective synthesis led to a paradigm shift and syntheses of stereochemically pure butyrolactones are also achieved.^41–49^ Recognizing the importance of deamination of amino acids to the corresponding substituted pyruvic acids, we undertook to synthesize isotetrones that possess variations of the substituent at *C*-4 carbon of the furanone scaffold.

A facile approach to deamination of L-amino acid is considered as an important strategy, in order to enable the incorporation of substituents at *C*-4 position of isotetrone.^50^ The reaction of L-amino acids **1** – **3**, isoleucine, phenylalanine and tyrosine, respectively, with trifluoroacetic anhydride, at optimal conditions of 85 °C and 1.5 h duration, afforded oxazolones **4** – **6** (**Scheme 1**), in good to moderate yields, in addition to the formation of trifluoroacetyl amino acids.^51,52^ Oxazolones **4** – **6** were subjected to aq. alkaline hydrolysis at room temperature for 18 h, the corresponding pyruvic acids **7** – **9**formed, possessing either keto- or the enol ether functionality.^53,54^ The enolization was observed higher in products **8** and **9**.

**Scheme 1.**
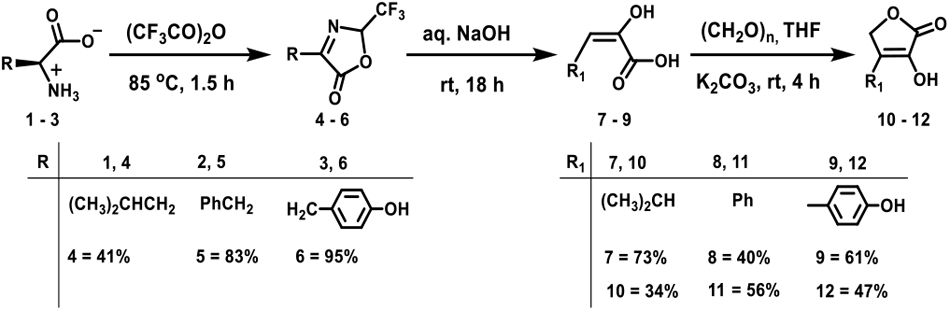
Synthesis of isotetrones **10** – **12**, from a-amino acids **1** – **3**, through oxazolones **4** – **6**and *C*-3 substituted pyruvic acids **7** – **9** intermediates.

The reaction of pyruvic acids**7** – **9** with formalin for 4 h, in the presence of K2CO3, in THF, initiated the aldolization and subsequent lactonization of the cross-aldol intermediate during acidic work-up led to the formation of isotetrones **10** – **12**. When the reaction was left for a longer duration, double aldolization product was also noticed in the crude reaction mixture, particularly, with pyruvic acid intermediate derived from isoleucine **1**. Whereas unreacted starting material **4** – **6** remained when aldolization reactions were conducted for a shorter duration. The identities and structural homogeneities of intermediates **4** – **9** and final products **10** – **12** were ascertained by NMR spectroscopies and mass spectrometry.

A similar reaction sequence was extended to valine, isoleucine and tryptophan amino acids. Oxozolone formation, hydrolysis to the corresponding pyruvic acid derivatives,^55–57^ aldolization with formalin, in the presence of K2CO3 and subsequent lactonization led to the formation of isotetrones **13** – **15**(**Fig. 1**). In the case of valine, intermediate a-keto-g-butyrolactone was observed to undergo a further reduction in the presence of formalin, so as to afford a-hydroxy-g-butyrolactone **13**, which is a pantolactone. Such a transformation was not observed with butyrolactone **14**, derived from isoleucine. Additional hydroxymethylation also occurred at the indole-nitrogen of tryptophan-derived butyrolactone **15**. Characterizations of products **13** – **15** were established by NMR spectroscopies and mass spectrometry.

**Figure 1.**
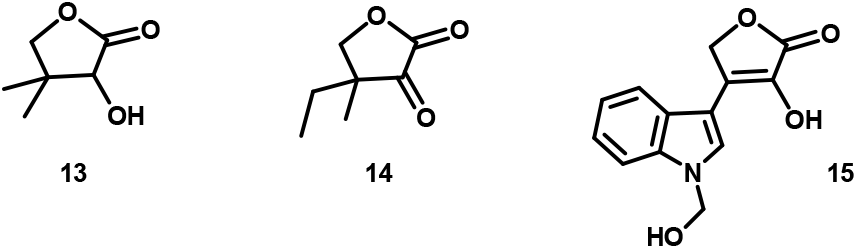
g-Butyrolactones **13** and **14**, and isotetronic acid **15** synthesized from valine, isoleucine and tryptophan, respectively.

### Specific isotetrones interfere with planktonic growth of M. smegmatis

The studies were undertaken with fast-growing *M. smegmatis* acid-fast bacterium. Of the six *C*-4 modified isotetrones **10** – **15**, preliminary screening of the effects on planktonic growth of the bacterium showed isotetrones **11** and **12** showed inhibitory properties and these derivatives were shortlisted for further studies. Isotetrone **11** showed significant growth inhibition at the early exponential and late exponential phases compared to the untreated controls (**Fig. 2a**).

**Figure 2.**
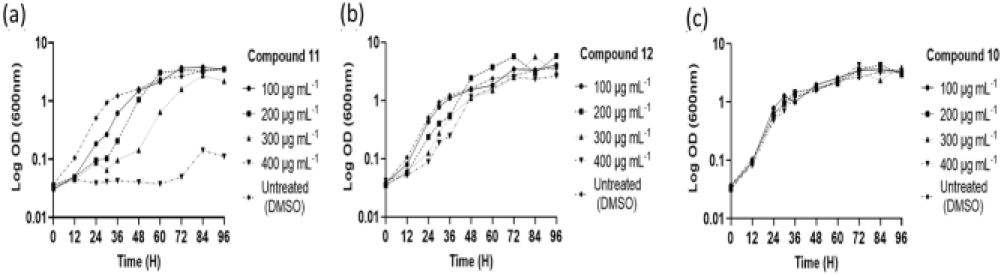
*M. smegmatis* growth curves in the presence of varying concentrations of (a) **11**; (b) **12** and (c) **10**.

Further analysis showed that derivative **11** inhibited the bacterial planktonic growth in a dose-dependent manner, and at 400 μg mL^-1^, complete growth inhibition occurred up to 72 h. After this duration, a re-growth of cells was observed, presumably due to the emergence of resistant mutants. A moderate growth inhibition was observed for isotetrone **12**, during the exponential growth phase of *M. smegmatis* cells in a concentration-dependent manner (Fig. 2b) and a maximal growth inhibition occurred at 400 μg mL^-1^. In this instance too, cells tended to recover from the early inhibition in the exponential phase and growth similar to untreated control was observed by 60-72 h, indicating a transient inhibitory effect of these isotetrones. Isotetrone **10** did not intervene in the growth profile of *M. smegmatis* cells across all concentrations when compared to the untreated control (Fig. 2c). These differences in the growth profiles indicate that both the isotetrones **11** and **12** possessing an aromatic moiety at *C*-4 carbon act as transient bacterial growth inhibitors.

### Isotetrones showed a differential killing pattern in M. smegmatis

The transient growth inhibition prompted us to verify the effects of isotetrones in combination with a known antimicrobial drug.^58,59^ A clinically relevant fluoroquinolone series drug, namely, ciprofloxacin (Cip), having a low minimum inhibitory concentration (MIC) value against *M. smegmatis*, was chosen for this purpose, which has been shown very effective and have a low MIC value against *M. smegmatis.* Initial determination of the MIC of Cip, in the presence of derivatives **10** – **12**(200 μg mL^-1^), showed the following trend: Cip: 0.25 μg mL^-1^; Cip +**10**: 0.25 μg mL^-1^; Cip + **11**: 0.125 μg mL^-1^ and Cip + **12**: 0.5 μg mL^-1^.

In order to assess the efficacies of isotetrones to kill *M. smegmatis* WT cells, in MB7H9 media, a kill kinetics assay was conducted, either alone or in combination with ciprofloxacin. Isotetrone derivatives alone did not result in any drop in CFU mL^-1^, implying that the derivatives lacked activity to kill the cells (**Fig. S13**). Whereas, an alternating killing pattern emerged when the cells were treated with a combination of ciprofloxacin (2.5 μg mL^-1^) and different isotetrones (400 μg mL^-1^). The drug in combination with **10** did not lead to any appreciable change in the killing, whereas ciprofloxacin (2.5 μg mL^-1^) and isotetrone **11** and **12** combinations led to an antagonistic response, where the combinations led to a reduced killing at each plating time point (**Fig. 3**). We presume the intrinsic growth inhibitory properties of **11** and **12** interfered with ciprofloxacin activity, leading to the differential killing of the replicating cells.

**Figure 3.**
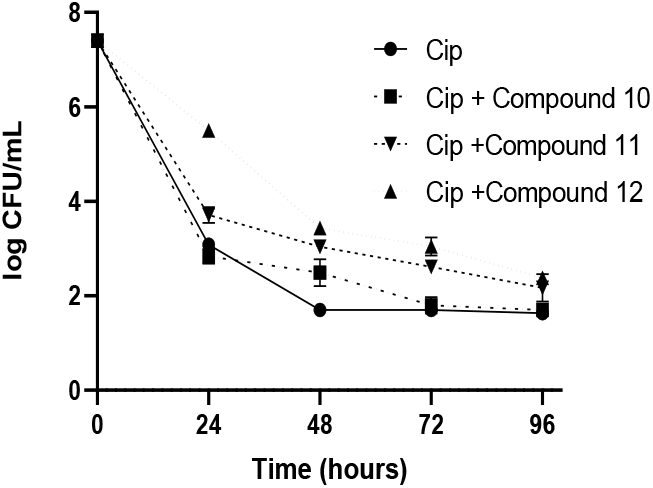
Kill kinetics of *M. smegmatis* with ciprofloxacin (Cip) (2.5 μg mL^-1^) and Cip in the presence of derivatives **10** – **12** (400 μg mL-^1^).

### Selective isotetrones affect the biofilm formation of M. smegmatis

Vitamin C is the first tetronic acid derivative demonstrated to exhibit inhibitory activity against biofilm-grown mycobacterium, occurring at a concentration of ~10 mM.^22^It was also assessed to interfere with the ppGpp biosynthesis in *M. smegmatis* and the biofilm growth.^60–63^ The isotetrones synthesized in the present work were assessed for the inhibition of biofilm formation at the maturation stage of *M*. *smegmatis*. The biofilm morphologies were monitored for 84 h post-exposure to **10** – **12**. Among the derivatives, **11** showed the largest effect in a concentration-dependent manner (**Fig. 4**). Whereas, derivative **12** showed moderate inhibition of the biofilm, derivative **10** did not show inhibition of biofilm, as that of the dense mature biofilm formed with untreated control (**Fig. 4**).

**Figure 4.**
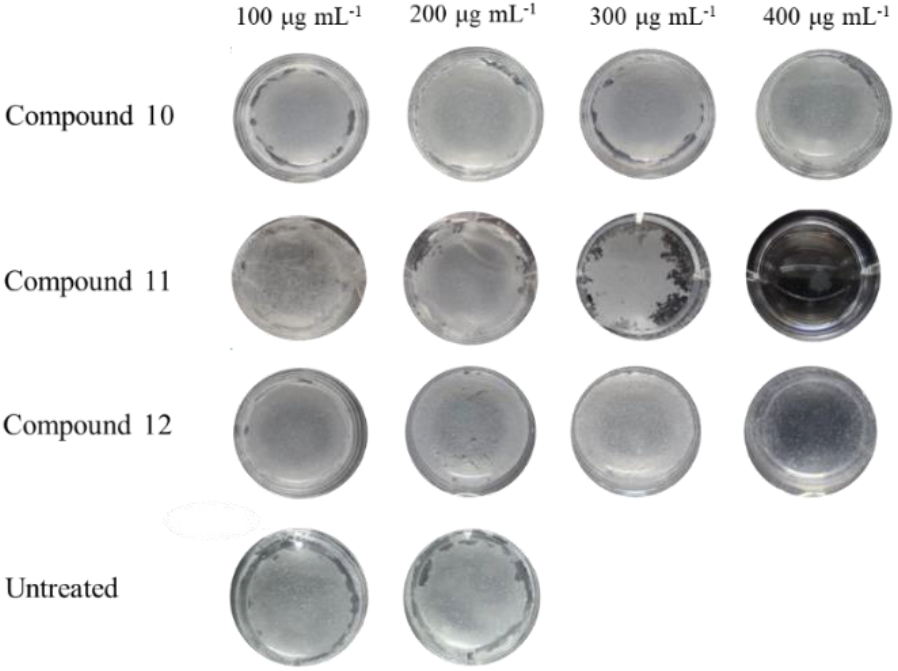
Images of *M. smegmatis* biofilms after 84 hours, in presence of varying concentrations of compounds **10** – **12** with untreated control.

### Isotetronic acid **12** intercepts with persister population regrowth and resistant mutant enrichment at low ciprofloxacin concentration

Earlier studies show that vitamin C targets the (p)ppGpp biosynthesis. The (p)ppGpp alarmone promotes the persister cell formation and plays a role in the antibiotic tolerance of the growing bacterium.^64–65^ Given this, further experiments were performed to identify whether selected isotetrones would intercept the persister population dynamics and resistant mutant enrichment at low ciprofloxacin concentration. In laboratory and clinical settings, antibiotic treatment often leads to a biphasic response, with a rapid killing phase followed by a plateau, which is represented by a 0.01-0.001% non-growing persister population growth. These antibiotic-tolerant persister cells lead to a major threat to effective antibiotic therapy, due to a resuscitation of the persisters into normal growing cells at a point soon after the antibiotic treatment is inadvertently terminated, which results in recurrent infection. It is thus of interest to identify the effect of the isotetronic acids, specifically that **11** and **12**, if these derivatives would possess a restrictive effect on the re-growth of the persister dominated antibiotic survived population, in presence of a lower concentration of ciprofloxacin (0.75 μg mL^-1^, which is 3 times higher than MIC). The lower concentration of the drug was chosen, as our data indicated the regrowth phenotype of drug-tolerant cells was very prominent at this concentration (**Fig. S14a**), as well as, genetic mutants enrichment (**Fig. S14b**). Further, the lower antibiotic concentration is very much relevant from a clinical perspective when optimum concentration could not be maintained. Next, the persister regrowth in presence of isotetrones **11** and **12** was checked. In this assay, cells were treated first with the drug (0.75 μg mL^-1^) for 24 h, thereby achieving a ~99.99% killing of the cells, leaving a residual ~0.01% persister population retained in the culture. In order to make sure the residual 0.01% after 24 h of ciprofloxacin treatment are bonafide persisters and not resistant mutants, the plating was done in MB+Cipro plates and hardly any resistant mutants were detected (data not shown). At this point, the culture was divided into 3 portions, added with either derivative **11** or **12** (400 μg mL^-1^) and an equal volume of sterile milliQ water. The growth kinetics was subsequently monitored, a brief non-growing phase of approximately 24 - 36 h was observed first and then antibiotic survived cells started to grow, as evident by the increase in CFU mL^-1^ at later time points. The data suggested significant prevention of the re-growth phase and a general delayed resuscitation of tolerant cells, in the presence of isotetrones **11** and **12** (400 μg mL^-1^). As shown in the resuscitation growth curve (**Fig. 5a**) between 48 and 72 h, the cells without isotetrones grew exponentially and reached a cell population comparable to that before the drug treatment was initiated. Whereas, in the presence of derivatives **11** and **12**, regrowth was prevented remarkably up to 72 h. Studies earlier reported on varied bacteria pointed out how persister cells could act as the predecessor of resistant mutants.^66–67^ In our assay since antibiotic survived cells were able to grow in presence of the same drug (ciprofloxacin), the possibility of emergence of resistant genetic mutants (cipro^R^) was estimated by plating parallelly in ciprofloxacin containing MB plates. A visible increase in cipro^R^ mutants was observed during the plateau/non-growing phase and, in the presence of isotetrones, a significant arrest of mutant enrichment occurred further. In the absence of isotetrones, namely, only in the presence of the drug, uninhibited resistant mutant enrichment over time occurred, with a remarkable ~98% mutant population taking over within 48 h of the regrowth phase. Both **11** and **12** derivatives significantly restricted resistant mutant enrichment for 48 h, presumably due to their inherent ability to slow re-growth of persisters and the subsequent transformation into mutants (**Fig. 5b**). Our data further suggested that the non-growing phase/plateau between 24 h and 48 h in the assay is responsible for conversion of persisters into genetic mutants and this critical step could be successfully prevented by isotetrones. These observations tempt us to raise and prove that isotetrones **11** and **12**, in combination with ciprofloxacin, act as bioenhancers and are therapeutically superior when administered together. The combination significantly inhibited re-growth of drug-tolerant population and arrested the resistant mutant enrichment. These findings implicate in the larger context of preventing resistance to antibiotics.

**Figure 5.**
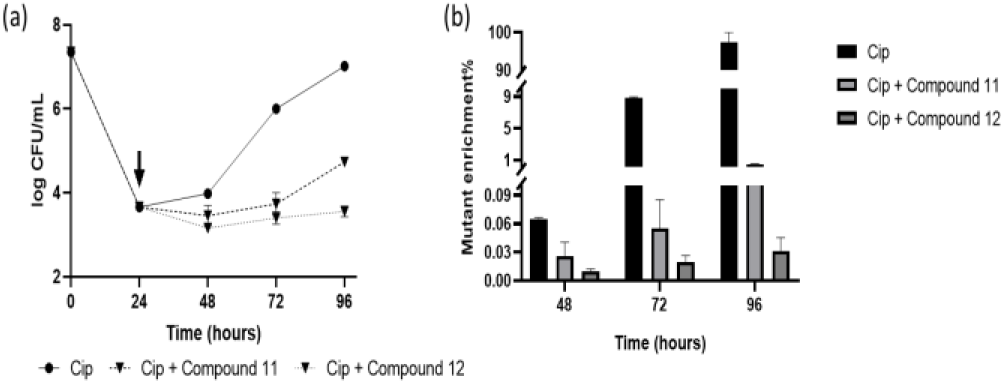
*M. smegmatis* delayed addition time-kill kinetics profile. (a) Reduction in the regrowth of ciprofloxacin (0.75 μg mL^-1^) treated tolearnt cells in the presence of compounds **11** and **12** (400 μg mL^-1^); (b) reduction of resistant mutant generation during ciprofloxacin (0.75μg mL^-1^) treatment in the presence of **11** and **12** (400 μg mL^-1^). Black arrows depict the time of compound addition.

### Isotetrone **12** inhibits long term starvation of cells under stringent conditions

In order to verify whether isotetrones would affect the survival kinetics under PBS starvation, cells in the PBS solution were incubated with the derivatives, at 400 μg mL^-1^ f.c. and CFU was estimated after 72 h for 2 weeks. The derivative **12** inhibited the growth and prevented long-term starvation in PBS (**Fig. 6**). The Long-term survival in PBS-starved conditions is linked to (p)ppGpp stress alarmone proliferation and earlier studies showed that vitamin C interfered with (p)ppGpp biosynthesis.^64–65^ Isotetrones **12** might possibly interfere with the stress alarmone biosynthesis similarly and thus a significant drop in CFU was observed even in the absence of an antibiotic under such stringent conditions. There are previous reports of vitamin C interacting with Rel enzyme^65^ and since our result also indicated a possible role of isotetrone **12** in regulating stringent response in *M. smegmatis*, we checked the possible interaction of compound **12** with RelMsm with molecular simulations.

**Figure 6.**
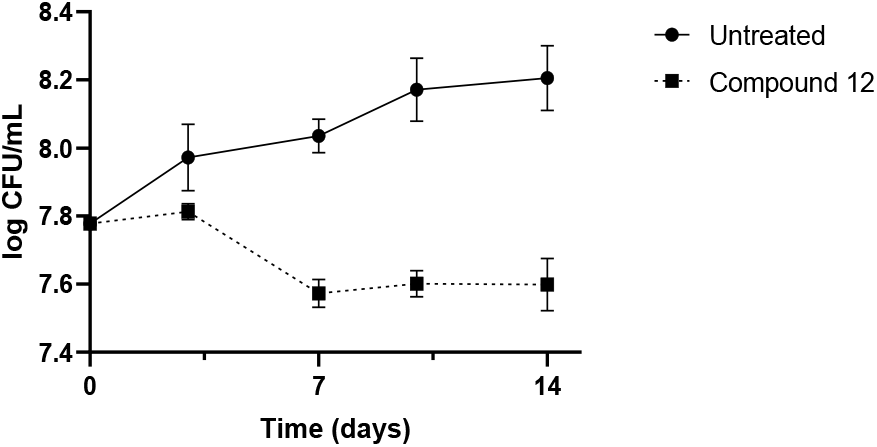
Plots of inhibition under long term starvation of *M. smegmatis* in PBS (untreated) and in the presence of compound **12.**

### Molecular dynamics simulation reveals notable interaction between compound 12 and Rel_Msm_

A systematic molecular dynamics (MD) simulation study was undertaken to assess the binding interaction of the isotetrone derivative **12** with Rel enzyme, with the knowledge that vitamin C binds directly to Rel enzyme and inhibits (p)ppGpp biosynthesis. **Figure 7** shows the Rel-ligand complex at their most energetically preferable binding sites, found from the docking study. The corresponding binding affinity is tabulated in **Table 2**. The structures served as initial configurations for subsequent MD simulation study. The protein-ligand complex structures after 10 ns of simulation are depicted in **Figure 8**. Note that, we run short 10 ns long MD simulation to get the most realistic binding site before calculating binding energy. The binding sites obtained in the docking study are preserved during the MD simulation. The average binding energies (ΔH_bind_) for ppGpp, compound **12** and vitamin C, obtained from the simulated trajectories, are presented in **Table 2**. It is evident that ppGpp binds to Rel protein most strongly, followed by compound **12** and vitamin C. The binding energy trend is in agreement with the docking results.

**Figure 7.**
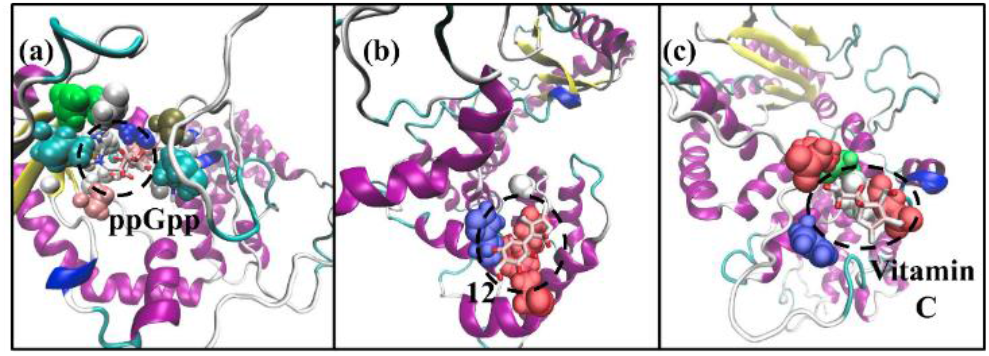
Protein-ligand complexes, obtained from docking studies. (a) Rel-ppGpp, (b) Rel-tyrosine lactone, and (c) relvitamin C complexes at their lowest energy conformations, predicted by molecular docking. These conformations served as the starting structures for MD simulation. The secondary structure of the protein has been represented with colour code: Yellow: β sheet, pale blue: turns, white: other residues. The ligands are presented using “licorice” representation scheme of VMD. Protein residues within a 5 Å distance from the ligands are presented using VdW spheres with a colour code: White: Non-polar residues, Blue: Basic residues, Red: Acidic residues, and Green: Polar residues. Water molecules are removed for clarity.

**Figure 8.**
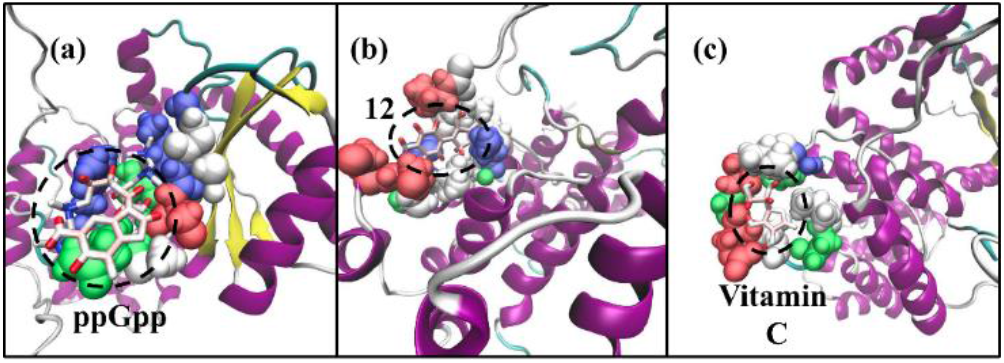
(a) Rel-ppGpp, (b) Rel-**12**, and (c) Rel-vitamin C complexes after 10 ns simulation. The secondary structure of the protein has been represented with colour code: Yellow: β sheet, pale blue: turns, white: other residues. The ligands are presented using “licorice” representation scheme of VMD. Protein residues within a 5 Å distance from the ligands are presented using VdW spheres with a colour code: White: Non-polar residues, Blue: Basic residues, Red: Acidic residues, and Green: Polar residues. Water molecules are removed for clarity.

**Table 2.** Binding affinity obtained from the docking study and average binding energy as calculated from MD-simulated trajectories using MM-GBSA.

| Ligand | $\Delta H(\text{kcal/mol})$<br>from Docking | $\Delta H(\text{kcal/mol})$<br>from MM-GBSA |
| --- | --- | --- |
| ppGpp | -7.23 | $-38.58 \pm 8.40$ |
| Compound **12** | -4.08 | $-9.74 \pm 3.37$ |
| Vitamin C | -2.91 | $-4.01 \pm 4.04$ |

**Figure 9a** presents the contributions of electrostatic and VdW components of the binding energy, for all the three ligands. For ppGpp, it can be easily seen that the protein-ligand binding is mostly driven by the electrostatic energy, whereas the contribution from the VdW interaction is not significant (**Fig. 9a**). This can be attributed to the presence of the phosphate groups on the ppGpp molecule. In the case of compound **12**, both the VdW and electrostatic energies significantly promoted the binding process (**Fig. 9a**). Like ppGpp, vitamin C also binds to Rel through coulombic interaction only, although the magnitude of the interaction was much smaller than the former, because of the absence of any charged functional groups. An interesting observation is that in case of vitamin C, the electrostatic contribution to ΔH is higher than that of compound **12**. Despite that, the binding affinity of vitamin C and Rel is weak, probably because of its higher solubility than compound **12**.

**Figure 9.**
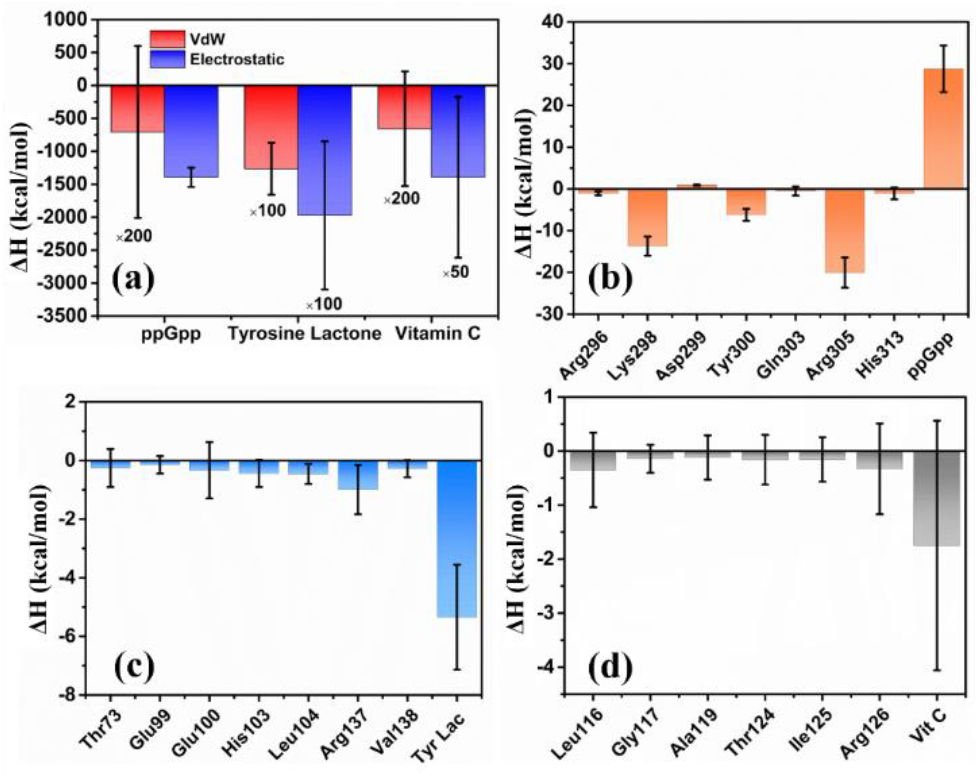
(a) Average electrostatic and VdW components of binding energy. Contribution of different residues in the binding energy for (b) Rel-ppGpp, (c) Rel-compound 12, and (d) Rel-vitamin C complexes. Nomenclature: Tyr Lac: compound 12; Vit C: vitamin C.

It is noteworthy that not all the residues of the Rel protein took part in the ligand binding phenomena. The per residue decomposition data of ΔG_bind_ is depicted in **Figures 9b-d**. In corroboration with the previous observations, the significant contribution towards Rel-ppGpp binding came from the positively charged residues like Arg and Lys (**Fig. 9b**). This is another evidence that the binding of ppGpp with Rel is dominated by the electrostatic interaction among the negatively charged phosphate groups present in ppGpp and the positively charged residues of the Rel. An interesting observation is that the contribution from ppGpp did not favour the protein-ligand binding (**Fig. 9b**) due to a high positive change in the polar solvation energy upon binding (1364.06 ±140.56 kcal/mol) to Rel, which indicates the presence of an attractive coulombic interaction between ppGpp and solvent molecules. For compound **12** and vitamin C, no specific protein residues provided a major contribution towards the binding energy, rather significant contribution came from the ligands (**Fig. 9c-d**). The absence of any charged groups in **12** and vitamin C can be attributed as the probable reason behind this striking difference.

Apart from the electrostatic and VdW interactions, proteinligand binding can also be mediated by the formation of hydrogen bonds (H-bonds) between the protein and ligand. We have used a cut off distance of 0.35 nm between donor-acceptor and a cut-off angle of 30° to define H-bonds. In case of ppGpp and Rel, Lys298, Tyr300, and Arg305 residues were involved in H-bonding. For compound 12, Glu100, His103, Arg137residues, and for vitamin C, Leu116, and Arg126 residues were found to form H-bonds with ligands. Average number of hydrogen bonds formed between ppGpp and Rel was found to be highest (9.23 ± 1.68), followed by **12** (1.18 ± 0.85), and vitamin C (1.31 ± 1.24). This occurred due to the presence of more H-bond forming acceptors and donors in ppGpp molecule, compared to compound **12** or vitamin C. Our simulation study confirms the experimental observation of the highest binding affinity of ppGpp, followed by compound **12** and vitamin C. The highest binding energy of ppGpp is attributed to the presence of the phosphate group and also to its ability to form more hydrogen bonds with Rel.

## Conclusion

The work herein demonstrates (i) chemical synthesis of hither-to unknown *C*-4 modified isotetrones from a-amino acids, through the formation of pyruvic acids, their aldonization and lactonization. (ii) Biochemical studies discover the role of these new isotetrones to interfere the growth of mycobacterium planktonic cells, inhibition of *M. smegmatis* biofilm formation is seen in a dose-dependent manner. (iii) Synergistic inhibitory effects are observed with well-known antibiotic ciprofloxacin in presence of the selected isotetronic acids, wherein the MIC of ciprofloxacin is improved by two-fold. (iv) Specific activities of isotetrones possessing aromatic substituents at *C*-4 carbon include the prevention of the re-growth of antibiotic tolerant population in *M. smegmatis*. We observe that *M. smegmatis* cells survive, form non-growing persisters and resuscitate in presence of lower concentrations of antibiotic ciprofloxacin, presumably due to enrichment of genetic mutants, as it occurs in a clinical scenario. (v) Systematic molecular dynamics simulations uncover the stabilities of Rel protein-inhibitor complexes, occurring through VdW, electrostatic and hydrogen-bonding interactions. The study shows that the addition of isotetrones hamper the regrowth stage of ciprofloxacin-selected persister population. Recurrent infection is a challenge in current antibiotic treatment regimens. The present work demonstrates the complete occlusion of the generation of resistant mutants, thereby opening up the feasibility to administer fewer concentrations of antibiotics and minimize associated off-target effects.

## Experimental section

### General procedure

Formalin and K_2_CO_3_ were sequentially added to the solution of keto-acid, derived from L-valine, L-leucine, L-isoleucine, L-phenylalanine, L-tyrosine and L-tryptophan) in THF, stirred at room temperature for 3 to 15 h. Tehe solution was evaporated *in vacuo* and the reaction mixture was treated with aq. HCl (3 N), extracted with Et_2_O (3 × 15 mL), organic portion washed with water (10 mL), brine (10 mL) and dried over Na_2_SO_4_ and concentrated *in vacuo*.

#### 3-Hydroxy-4-isopropylfuran-2(5*H*)-one (10)

Formalin (0.38 mL, 4.62 mmol) and K_2_CO_3_ (0.66 g, 4.84 mmol) were sequentially added to a stirred solution of **7**^53^ (0.5 g, 3.85 mmol) in THF (5 mL), stirred for 4 h at room temperature and worked up as described in the general procedure. The crude product was purified to afford **10** (0.12 g, 34%), as a colourless oil. Rf (pet. ether/EtOAc = 7:3) 0.4. ^1^H NMR (CDCl_3_, 400 MHz): δ 4.68 (s, 2 H, C*H*_2_O), 2.89 (septet, *J* = 20.8, 13.6, 6.8 Hz, 1 H, C*H*(CH_3_)_2_), 1.18 (d, *J* = 6.8 Hz, 6 H, CH(C*H*_3_)_2_); ^13^C NMR (CDCl_3_, 100 MHz): δ 171.7, 137.5, 135.6, 66.1, 25.5, 20.4. GCMS: *m/z* calcd. for C_7_H_10_O_3_: 142 [M+], found 142, with fragments at *m/z* at 97, 69 and 41.

#### 3-Hydroxy-4-phenylfuran-2(5*H*)-one (11)

Formalin (0.09 mL, 1.08 mmol), K_2_CO_3_ (0.14 g, 1.14 mmol) were added sequentially to solution of **8**^54^ (0.15 g, 0.9 mmol) in THF (2.5 mL), the reaction mixture stirred for 4 h at room temperature and worked-up as described in the general procedure. The residue was purified to afford **11** (0.073 g, 56%), as a white solid. R_f_ (pet. ether/EtOAc = 9:1) 0.7. ^1^H NMR (CD_3_OD, 400 MHz): δ 7.66 (d, *J*= 4 Hz, 2 H, C_6_*H*_5_), 7.36 (t, *J* = 8 Hz, 2 H, C_6_*H*_5_), 7.29-7.26 (m, 1 H, C_6_*H*_5_), 5.03 (s, 2H, C*H*_2_O); ^13^C NMR (CD_3_OD, 400 MHz): δ 172.3, 138.8, 134, 132.2, 130.7, 129.9, 129.7, 129.4, 127.6, 69.2. HRMS: m/z calcd. for C_10_H_8_O_3_: 177.0552[M+H]^+^, found 177.0551.

#### 3-Hydroxy-4-(4-hydroxyphenyl)furan-2(5*H*)-one (12)

Formalin (0.16 mL, 1.94 mmol) and K_2_CO_3_ (0.28 g, 2.04 mmol) were successively added to a solution of **9**^54^ (0.35 g, 1.94 mmol) in THF (5 mL), stirred for 4 h at room temperature and worked up as described in the general procedure. Column chromatography purification of the residue afforded **12** (0.127 g, 47%), as a white solid. R_f_ (pet. ether/EtOAc = 3:2) 0.5. ^1^H NMR (CD_3_OD, 400 MHz): δ 7.59 (d, *J* = 8.8 Hz, 2 H, C_6_*H*_5_), 6.83 (d, *J* = 8.4 Hz, 2 H, C_6_*H*_5_), 5.06 (s, 2 H, C*H*_2_O); ^13^C NMR (CD_3_OD, 100 MHz): δ 172.8, 159.6, 136.5, 129.3, 128.8, 123.6, 116.6, 69.2. HRMS: m/z calcd. for C_10_H_9_O_4_: 193.0501[M+H]^+^, found 193.0501.

#### 3-Hydroxy-4,4-dimethyldihydrofuran-2-one (13)

Formalin (1.75 mL, 21.5 mmol) and K_2_CO_3_ (1.5 g, 10.75 mmol) were added successively to a solution of 2-oxoisovaleric acid^55^ (0.5 g, 4.3 mmol) in THF (5 mL), the reaction was stirred for 15 h at room temperature and worked up as described in the general procedure. Vacuum distillation was performed to remove the unreacted starting material and column chromatography purification of the residue gave **13** (0.19 g, 34%) as a white solid. R_f_ (pet. ether/EtOAc = 9:1) 0.8. ^1^H NMR (CDCl_3_, 400 MHz): δ 4.15 (s, 1 H, C*H*OH), 3.98 (d, *J* =9.2 Hz, 1 H, C*H*_2_O), 3.92 (d, *J* = 9.2 Hz, 1 H, C*H*_2_O), 1.18 (s, 3 H, *CH_3_*), 1.03 (s, 3 H, *CH_3_*);^13^C NMR (CDCl3, 100 MHz): δ 178.1, 76.4, 75.6, 40.7, 22.6, 18.7. GCMS: *m/z* calcd. for C_6_H_10_O_3_: 131 [M+H], found 131, with fragments at *m/z* at 71, 57, 43 and 41.

#### 4-Ethyl-4-methyl dihydrofuran-2,3-dione (14)

Formalin (1.9 mL, 23.1 mmol) and K_2_CO_3_ (1.6 g, 11.55 mmol) were added successively to a solution of 3-methyl-2-oxopentanoic acid^57^ (0.5 g, 3.85 mmol) in THF (5 mL), stirred for 15 h and worked-up as described in the general procedure. The residue was purified by vacuum distillation to afford **14** (0.29 g, 54%), as a yellow liquid. (R_f_ (pet. ether/EtOAc = 9:1) 0.8. ^1^H NMR (CDCl_3_, 400 MHz): δ 4.55 (d, *J* = 9.6 Hz, 1 H, C*H*_2_O), 4.39 (d, *J* = 9.6 Hz, 1 H, C*H*_2_O), 1.78-1.62 (m, 2 H, C*H*_2_CH_3_), 1.26 (s, 3 H, C*H*_3_), 0.92 (t, *J* = 7.2 Hz, 3 H, CH_2_C*H*_3_); ^13^C NMR (CDCl_3_, 100 MHz): δ 198.4, 161, 75.6, 45.4, 29.3, 19.4, 8.2, 8. GCMS: *m/z* calcd. for C_7_H_10_O_3_: 142 [M+], found 142, with fragments at *m/z* at 85, 71, 69, 55 and 41.

#### 3-Hydroxy-4-(1-hydroxymethyl)-1*H*-indol-3-yl)furan-2(5*H*)-one (15)

Formalin (0.22 mL, 2.71 mmol) and K_2_CO_3_ (0.38 g, 2.71 mmol) were added successively to a solution of indole-3-pyruvic acid^56^ (0.5 g, 2.46 mmol) in THF (5 mL) was treated with, and the reaction was stirred for 6 h at room temperature and worked up as described in the general procedure. The crude product was purified to afford **15** (0.174 g, 33%), as a white solid. R_f_ (pet. ether/EtOAc = 3:1) 0.3. ^1^H NMR (CD_3_OD, 400 MHz): δ 7.96 (s, 1 H, C*H*NH), 7.82 (d, *J* = 8 Hz, 1 H, C_6_*H*_5_), 7.59 (d, *J* = 8 Hz, 1 H, C_6_*H*_5_), 7.28 (t, *J* = 8 Hz, 1 H, C_6_*H*_5_), 7.19 (t, *J* = 8 Hz, 1 H, C_6_*H*_5_), 5.63 (s, 2 H, C*H*_2_O), 5.29 (s, 2 H, C*H*_2_O); ^13^C NMR (CD_3_OD, 100 MHz): δ 171.2, 136.1,133.1, 129.2, 126.4, 126.2, 122.3, 120.7, 120.5, 110.1, 107, 69, 68.1. HRMS: m/z calcd. for C_13_H_11_NO_4_: 246.0766[M+H]^+^, found 246.0767.

### Bacterial growth and culture conditions

For all the assays, *M. smegmatis* mc^2^155 strain was grown in MB7H9 media (HiMedia) containing 0.05% tween-80 and 2% glucose; agar (1.6%, w/v) (Himedia) was used to make agar plates. Ciprofloxacin powder was obtained from Sisco Research Laboratories. Antibiotics were used at variable concentrations. Unless mentioned otherwise, strains were grown at 37 °C and 150 rpm.

### Growth inhibition assay

For analyzing the effect of isotetrones on the growth of the *M. smegmatis* mc^2^155 strain, bacteria were grown till the mid-exponential phase, further inoculation was done in MB7H9 media containing 0.05% tween-80 and 2% glucose to make the final OD ~ 0.03. 3 mL of such culture was taken in a sterile glass tube and 30 μL of each of the respective isotetrones with varying final concentrations (100 μg mL^-1^, 200 μg mL^-1^, 300 μg mL^-1^, and 400 μg mL^-1^) were added to the media at 0 hours and OD at 600 nm was measured. The bacterial growth was further monitored in equal intervals for 96 h by recording OD600 and plotted using Graph Pad Prism Software. The tube containing no compounds (only DMSO) served as the untreated positive control to observe and compare growth inhibition in the presence and absence of the compounds. This experiment was performed in a set of 2 experimental replicates.

### Biofilm formation assay

The primary culture for the wildtype strain *M. smegmatis* mc^2^ 155 was grown in MB7H9 medium at 37 °C, 150 rpm shaking. The cells were harvested at the stationary phase and washed with Sauton’s media (Himedia) twice, the cell pellet was resuspended in the sauton’s media containing 2% glucose and the final OD was adjusted to 0.05. Subsequently, 200 μL of culture was poured into a well of sterile 96 well microtiter plates supplemented with the 2 μL respective isotetrones having a final concentration of 100 μg mL^-1^, 200 μg mL^-1^, 300 μg mL^-1^, and 400 μg mL^-1^. The control wells included in the same plate were untreated control (only cells, no compounds to see uninhibited biofilm formation) and media control (only media, no cells to check contamination). The inoculated plates were sealed and kept in a 37 °C humidified incubator to avoid drying of media and incubated for a minimum of 72-96 h without any external disturbance. The images were captured under white light at different time points. The experiment was performed in a set of 3 biological replicates.

### Minimum inhibition concentration (MIC) determination

For analyzing if the isotetrones could increase the susceptibility to ciprofloxacin, we performed a MIC assay using the ciprofloxacin in combination with compounds (200 μg mL^-1^ f.c.) by broth microdilution assay using resazurin dye. ciprofloxacin was 2-fold diluted in the 96-well microplate to achieve a broad range of final concentrations ranging from 16 μg mL^-1^ to 0.03 μg mL^-1^. *M. smegmatis* Cultures were grown till mid-exponential phase (OD_600_ of 0.6-1), diluted to OD_600_ of 0.01 in fresh MB7H9 medium, and 196 μL were added in each well of the 96-well plate containing 4 μL of different concentrations of ciprofloxacin. Depending on the series, cells were pre-mixed with different isotetrones (200 μg mL^-1^ f.c.), in the no-compound series equal amount of sterile miliQ was added. Following the incubation of 36 h standing at 37 °C, 30 μL of 0.1 mg mL^-1^ of resazurin was added to each of the wells and again incubated for 6 to 8 h at 37 °C, and dye reduction value was recorded.

### Time-kill kinetics assay

*M. smegmatis* cultures were grown till the late exponential/stationary phase (OD_600_ of 1.5-2.5), diluted 1:100 to start a secondary culture and grown till the mid-exponential phase (OD_600_ of 0.6-1), and further diluted to adjust to OD_600_ of 0.2 corresponding to ~2 ×10^7^ CFU mL^-1^ (CFU= colony-forming unit) in fresh MB7H9 medium. 3 mL of such culture was added to one tube and 2.5 μg mL^-1^ (f.c.) ciprofloxacin (10X MIC) was added with or without isotetrones (400 μg mL^-1^) at T-0 and CFU mL^-1^. was estimated by spreading into the MB7H9 agar plate. Tubes were incubated at 37 °C shaker and CFU estimations (plating) were done every 24 h up to 96 h. Plates were further incubated at 37 °C for 3-4 days for colony growth and subsequently counted. To measure the killing efficiency of the compounds alone (without ciprofloxacin) similar kill kinetics assay was set up in parallel. All Kill kinetics assays were performed with at least 3 biological replicates.

### Delayed addition of isotetrones in time-kill kinetics

*M. smegmatis* secondary culture was prepared as mentioned above and then 0.75 μg mL^-1^ of ciprofloxacin (3X MIC) was added to the media. After 24 h of incubation, the culture was divided into separate tubes and different isotetroneswere added (400 μg mL^-1^ f.c.) into it. CFU estimations (plating) were done every 24 h up to 96 h. The tube containing no compounds (only DMSO) served as the untreated control to compare regrowth in absence of the compounds. This experiment was performed in a set of 3 replicates.

### Mutant enrichment determination

In parallel to the CFU estimation in Kill kinetics experiments, spotting/spreading was done in ciprofloxacin (1.25 μg mL^-1^) plates to enumerate resistant mutant population enrichment over time.

### PBS starvation assay

*M. smegmatis* cultures were grown till the late exponential/stationary phase (OD_600_ of 1.5-2.5), diluted 1:100 to start a secondary culture, and grow till the mid-exponential phase (OD_600_ of 0.6-1). After that, the cells were washed twice with PBS and resuspended in an equal volume of PBS. T-0 CFU mL^-1^. estimation was done immediately and the culture was divided into equal volumes in several glass tubes followed by the addition of selective isotetrones (400 μg mL^-1^. f.c.). In the untreated control tube, an equal volume of sterile MilliQ water was added. CFU estimation was done every 72 h for ~ 3 weeks.

## Supporting Information

Docking and molecular dynamics simulation protocol, NMR Spectra of new compounds, Kill kinetics and persister regrowth.

## AUTHOR INFORMATION

### Author contributions

D.C., P.M., A.G. and N.J. contributed to the conception and design of the study. K.B., A.P., S.B. M.S. and A.G. performed the experiments. D.C., P.M., A.G. and N.J. participated in data analysis and interpretation. All authors participated in writing the manuscript.

## Acknowledgment

Authors thank the Department of Biotechnology (DBT), Government of India, for funding this work (Grant number: BT/PR33123/MED/29/1497/2020). Indian Institute of Science, Bangalore, is gratefully acknowledged for a research fellowship to K.B.

## REFERENCES

(1) Harms, A.; Maisonneuve, E.; Gerdes, K. Mechanisms of Bacterial Persistence During Stress and Antibiotic Exposure. Science 2016, 354, aaf4268.

(2) Chatterji, D.; Ojha, A. K. Revisiting the Stringent Response, ppGpp And Starvation Signaling. Curr. Opin. Microbiol. 2001, 4, 160–165.

(3) Srivatsan, A.; Wang, J. D. Control of Bacterial Transcription, Translation and Replication by (p)ppGpp. Curr. Opin. Microbiol. 2008, 11, 100–105.

(4) Hengge, R., “High-Specificity Local and Global c-di-GMP Signaling”, Trends Microbiol. 2021, 29, 993–1003.

(5) Kushwaha, G. S.; Oyeyemi, B. F.; Bhavesh, N. S. Stringent Response Protein as A Potential Target to Intervene Persistent Bacterial Infection. Biochimie 2019, 165, 67–75.

(6) Sharma, I. M.; Petchiappan, A.; Chatterji, D. Quorum Sensing and Biofilm Formation in Mycobacteria: Role of c-di-GMP and Methods to Study This Second Messenger. IUBMB Life 2014, 66, 823–834.

(7) Hogg, T.; Mechold, U.; Malke, H.; Cashel, M.; Hilgenfeld, R. Conformational Antagonism Between Opposing Active Sites in A Bifunctional RelA/SpoT Homolog Modulates (p)ppGpp Metabolism During the Stringent Response [corrected]. Cell 2004, 117, 57–68.

(8) Page, R.; Peti, W. Toxin-Antitoxin Systems in Bacterial Growth Arrest and Persistence. Nat. Chem. Biol. 2016, 12, 208–214.

(9) Syal, K.; Joshi, H.; Chatterji, D.; Jain, V. Novel pppGpp Binding Site at the C-Terminal Region of the Rel Enzyme from *Mycobacterium smegmatis*. FEBS J. 2015, 282, 3773–3785.

(10) Liu, S.; Wu, N.; Zhang, S. S.; Yuan, Y. H.; Zhang, W. H. Zhang, Y. Variable Persister Gene Interactions with (p)ppgpp for Persister Formation in *Escherichia coli*. Front Microbiol. 2017, 8, 1795.

(11) Tian, C.; Semsey, S.; Mitarai, N. Synchronized Switching of Multiple Toxin–Antitoxin Modules by (p)ppGpp Fluctuation. Nucleic Acids Res. 2017, 45, 8180–8189.

(12) Tkachenko, A. G.; Kashevarova, N. M.; Sidorov, R.Y.; Shipilovskikh, S. A.; Rubtsov, A. E.; Malkov, A. V. A Synthetic Diterpene Analogue Inhibits Mycobacterial Persistence and Biofilm Formation by Targeting (p)ppGpp Synthetases. Cell Chemical Biology 2021, 28, 1420–1432.

(13) Rhee, H. W.; Lee, C. R.; Cho, S. H.; Song, M. R.; Cashel, M.; Choy, H. E.; Seok, Y. J.; Hong, J. I. Selective Fluorescent Chemosensor for the Bacterial Alarmone (p)ppGpp. J. Am. Chem. Soc. 2008, 130, 784–785.

(14) Wexselblatt, E.; Kaspy, I.; Glaser, G.; Katzhendler, J.; Yavin, E. Design, Synthesis and Structure-Activity Relationship of Novel Relacin Analogs as Inhibitors of Rel Proteins. Eur. J. Med. Chem. 2013, 70, 497–504.

(15) Sureka, K.; Ghosh, B.; Dasgupta, A.; Basu, J.; Kundu, M.; Bose, I. Positive Feedback and Noise Activate the Stringent Response Regulator Rel in Mycobacteria. PLoS One 2008, 3, e1771.

(16) Syal, K.; Bhardwaj, N.; Chatterji, D. Vitamin C Targets (p)ppGpp Synthesis Leading to Stalling of Long-Term Survival and Biofilm Formation in *Mycobacterium smegmatis*. FEMS Microbiol. Lett. 2017, 364, fnw282 (1–6).

(17) Vilchèze, C.; Kim, J.; Jacobs, Jr. W. R. Vitamin-C Potentiates the Killing of *Mycobacterium tuberculosis* by the First-Line Tuberculosis Drugs Isoniazid and Rifampin in Mice. Antimicrob. Agents Chemother. 2018, 62, e02165–17.

(18) Vilchèze, C.; Hartman, T.; Weinrick, B.; Jacobs, Jr. W. R. *Mycobacterium tuberculosis* is Extraordinarily Sensitive to Killing by A Vitamin C-Induced Fenton Reaction. Nat. Commun. 2013, 4, 1881.

(19) Keren, I.; Kaldalu, N.; Spoering, A.; Wang, Y.; Lewis, K. Persister Cells and Tolerance to Antimicrobials. FEMS Microbiol. Lett, 2004, 230, 13–18.

(20) Peddireddy, V.; Doddam, S. N.; Ahmed, N. Mycobacterial Dormancy Systems and Host Responses in Tuberculosis. Front Immunol., 2017, 8(84), 1–19.

(21) T, J. A. S.; J, R.; Rajan, A.; Shankar, V. Features of the Biochemistry of *Mycobacterium smegmatis,* as a Possible Model for *Mycobacterium tuberculosis*. J. Infect. Public Health 2020, 13, 1255–1264.

(22) Princiotto, S.; Jayasinghe, L.; Dallavalle, S. Recent Advances in the Synthesis of Naturally Occurring Tetronic Acids. Bioorganic Chem. 2022, 119, 105552 (1 – 32).

(23) Fischer, P. M.; Lane, D. P. Inhibitors of Cyclin-Dependent Kinases as Anti-Cancer Therapeutics. Current Med. Chem. 2000, 7,1213 – 1245.

(24) Nishio, K.; Ishida, A.; Arioka, H.; Kurokawa, H.; Fukuoka, K.; Nomoto, T.; Fukumoto, H.; Yokote, H.; Saijo, N. Antitumor Effects of Butyrolactone I, A Selective cdc2 Kinase Inhibitor, on Human Lung Cancer Cell Lines. Anticancer Res. 1996, 16, 3387–3395.

(25) Suzuki, M.; Hosaka, Y.; Matsushima, H.; Goto, T.; Kitamura, T.; Kawabe, K. Butyrolactone I Induces Cyclin B1 and Causes G2/M Arrest and Skipping of Mitosis in Human Prostate Cell Lines. Cancer Lett. 1999, 138, 121 – 130.

(26) Parvatkar, R. R.; D’Souza, C.; Tripathi, A.; Naik, C. G. Aspernolides A and B, Butenolides from A Marine-Derived Fungus *Aspergillus terreus*. Phytochemistry 2009, 70, 128–132.

(27) Haritakun, R.; Rachtawee, P.; Chanthaket, R.; Boonyuen N.; Isaka, M. Butyrolactones from the Fungus *Aspergillus terreus* BCC 4651. Chem. Pharm. Bull. 2010, 58, 1545 – 1548.

(28) Adpressa D. A.; Loesgen, S. Bioprospecting Chemical Diversity and Bioactivity in A Marine Derived *Aspergillus terreus*. Chem. Biodiversity 2016, 13, 253 – 259.

(29) Dewi, R. T.; Tachibana, S.; Darmawan, A. Effect On A-Glucosidase Inhibition and Antioxidant Activities of Butyrolactone Derivatives from *Aspergillus terreus* MC751. Med. Chem. Res. 2014, 23, 454 – 460.

(30) Niu, X.; Dahse, H. M.; Menzel, K. D.; Lozach, O.; Walther, G.; Meijer, J.; Grabley, S.; Sattler, I. Butyrolactone I Derivatives from *Aspergillus terreus* Carrying an Unusual Sulfate Moiety. J. Nat. Prod. 2008, 71, 689 – 692.

(31) Sugiyama, Y.; Yoshida, K.; Abe, N.; Hirota, A. Soybean Lipoxygenase Inhibitory and DPPH Radical-Scavenging Activities of Aspernolide A and Butyrolactones I and II. Biosci. Biotechnol. Biochem. 2010, 74, 881 – 883.

(32) Brachmann, A. O.; Forst, S.; Furgani, G. M.; Fodor, A.; Bode, H. B. Xenofuranones A and B: Phenylpyruvate Dimers from *Xenorhabdus szentirmaii*. J. Nat. Prod. 2006, 69, 1830 – 1832.

(33) Ingerl, A.; Justus, K.; Hellwig, V.; Steglich, W. Syntheses of Retipolide E and Ornatipolide, 14-Membered Biaryl-ether Macrolactones from Mushrooms. Tetrahedron 2007, 63, 6548 – 6557.

(34) Justus, K.; Herrmann, R.; Klamann, J.-D.; Gruber, G.; Hellwig, V.; Ingerl, A.; Polborn, K.; Steffan B.; Steglich, W. Retipolides - Unusual Spiromacrolactones from the Mushrooms *Retiboletus retipes* and *R. ornatipes*. Eur. J. Org. Chem. 2007, 5560–5572.

(35) Nitta, K.; Fujita, N.; Yoshimura, T.; Arai, K.; Yamamoto, Y. Metabolic Products of *Aspergillus Terreus*. IX. Biosynthesis of Butyrolactone Derivatives Isolated from Strains IFO 8835 and 4100. Chem. Pharm. Bull. 1983, 31, 1528 – 1533.

(36) Machajewski, T. M.; Wong, C.-H. The Catalytic Asymmetric Aldol Reaction. Angew. Chem. Int. Ed. 2000, 39, 1352 – 1374.

(37) Allen, S. T.; Heintzelman, G. R.; Toone, E. J. Pyruvate Aldolases as Reagents for Stereospecific Aldol Condensation. J. Org. Chem. 1992, 57, 426–427.

(38) Enders, D.; Dyker, H.; Leusink, F. R. Enantioselective Synthesis of Protected Isotetronic Acids. Chem. Eur. J. 1998, 4, 311 – 320.

(39) Dambruoso, P.; Massi, A.; Dondoni, A. Efficiency in Isotetronic Acid Synthesis Via A Diamine-acid Couple Catalyzed Ethyl Pyruvate Homoaldol Reaction. Org. Lett. 2005, 7, 4657 – 4660.

(40) Zhou, Z.; Walleser, P. M.; Tius, M. A. Isotetronic Acids from an Oxidative Cyclization. Chem. Commun. 2015, 51, 10858 – 10860.

(41) Mao, B.; Fan anas-Mastral, M.; Feringa, B. L. Catalytic Asymmetric Synthesis of Butenolides and Butyrolactones. Chem. Rev. 2017, 117, 10502 – 10566.

(42) Juhl, K.; Gathergood, N.; Jørgensen, K. A. Catalytic Asymmetric Homo-Aldol Reaction of Pyruvate-A Chiral Lewis Acid Catalyst that Mimics Aldolase Enzymes. Chem. Commun. 2000, 2211 – 2212.

(43) Gathergood, N.; Juhl, K.; Poulsen, T. B.; Thordrup, K.; Jørgensen, K. A. Direct Catalytic Asymmetric Aldol Reactions of Pyruvates: Scope and Mechanism. Org. Biomol. Chem. 2004, 2, 1077 – 1085.

(44) Vincet, J. M.; Margottin, C.; Berlande, M.; Cavagnat, D.; Buf-feteau, T.; Landais, Y. A Concise Organocatalytic and Enantiose-lective Synthesis of Isotetronic Acids. Chem. Commun. 2007, 4782 – 4784.

(45) Roy, B.; Das, E.; Roy, A.; Mal, D. Ni(ii)-Catalyzed Vinylic CH Functionalization of 2-Acetamido-3-Arylacrylates to Access Isotetronic Acids. Org. Biomol. Chem. 2020, 18, 3697 – 3706.

(46) Chen, P.; Wang, K.; Zhang, B.; Guo, W.; Liu, Y.; Li, C. Water Enables an Asymmetric Cross Reaction of α-Keto Acids With α-Keto Esters for the Synthesis of Quaternary Isotetronic Acids. Chem. Commun. 2019, 55, 12813 – 12816.

(47) Zhang, B.; Jiang, Z.; Zhou, X.; Lu, S.; Li, J.; Liu, Y.; Li, C. The Synthesis of Chiral Isotetronic Acids with Amphiphilic Imidazole/ Pyrrolidine Catalysts Assembled in Oil-In-Water Emulsion Droplets. Angew. Chem., Int. Ed. 2012, 51, 13159 – 13162.

(48) Xu, X.-Y.; Tang, Z.; Wang, Y.-Z.; Luo, S. W.; Cun, L. F.; Gong, L.-Z. Asymmetric Organocatalytic Direct Aldol Reactions of Ketones with α-keto acids and their Application to the Synthesis of 2-Hydroxy-γ-Butyrolactones. J. Org. Chem. 2007, 72, 9905 – 9913.

(49) Lee, D.; Newman, S. G.; Taylor, M. S. Boron-Catalyzed Direct Aldol Reactions of Pyruvic Acids. Org. Lett. 2009, 11, 5486 – 5489.

(50) Cooper, A. J. L.; Ginos, J. Z.; Meister, A. Synthesis and Properties of the α-Keto Acids. Chem. Rev. 1983, 83, 321–358.

(51) Liu, Y.-X.; Zhang, P.-X.; Li, Y.-Q.; Song, H.-B.; Wang, Q.-M. Design, Synthesis, and Biological Evaluation Of 2-Benzylpyrroles and 2-Benzoylpyrroles Based on Structures of Insecticidal Chlorfenapyr and Natural Pyrrolomycins. Molecular Diversity. 2014, 18, 593–598.

(52) Gräßle, S.; Vanderheiden, S.; Hodapp, P.; Bulat, B.; Nieger, M.; Jung, N.; Bräse, S. Solid Phase Synthesis of (Benzannelated) Six-Membered Heterocycles via Cyclative Cleavage of Resin-Bound Pseudo-Oxazolones. Org. Lett. 2016, 18, 3598–3601.

(53) Furukawa, K.; Inada, H.; Shibuya, M.; Yamamoto, Y. Chemoselective Conversion from α-Hydroxy Acids to α-Keto Acids Enabled by Nitroxyl-Radical-Catalyzed Aerobic Oxidation. Org. Lett. 2016, 18, 4230–4233.

(54) Yu, J.; Li, J.; Cao, S.; Wu, T.; Zeng, S.; Zhang, H.; Liu, J.; Jiao, Q. Chemoenzymatic Synthesis of L-3,4-Dimethoxyphenyl-Alanine and Its Analogues Using Aspartate Aminotransferase as A Key Catalyst. Catalysis Comm. 2019, 120, 28–32.

(55) Endo, Y.; Shudo, K.; Itai, A.; IIasegawa, M.; Sakai, S. Synthesis and Stereochemistry of Indolactam-V, An Active Fragment of Teleocidins. Structural Requirements for Tumor-Promoting Activity. Tetrahedron 1986, 42(21), 5905–5924.

(56) Bergman, J.; Lidgren, G.; Gogoll, A. Synthesis and Reactions of Oxazolones from L-Tryptophan and α-Haloacetic Anhydrides. Bull. Soc. Chim. Belg. (European Section) 1992, 101(7), 643–660.

(57) Lin, D.W.; Masuda, T.; Biskup, M.B.; Nelson, J.D.; Baran, P.S. Synthesis-Guided Structure Revision of the Sarcodonin, Sarcoviolin, and Hydnellin Natural Product Family. J. Org. Chem. 2011, 76, 1013–1030.

(58) Sikri, K.; Duggal, P.; Kumar, C.; Batra, S. D.; Vashist, A.; Bhaskar, A.; Tripathi, K.; Sethi, T.; Singh, A.; Tyagi, J. S. Multifaceted Remodeling by Vitamin C Boosts Sensitivity of *Mycobacterium Tuberculosis* Subpopulations to Combination Treatment by Anti-Tubercular Drugs. Redox Biol. 2018, 5, 452 – 466.

(59) Khameneh, B.; Bazzaz, B. S. F.; Amani, A.; Rostami, J.; Vahdati-Mashhadian, N. Combination of Anti-Tuberculosis Drugs with Vitamin C or NAC Against Different *Staphylococcus Aureus* and *Mycobacterium Tuberculosis* Strains. Microb. Pathog. 2016, 93, 83–87.

(60) Vilcheze, C., Hartman, T., Weinrick, B., and Jacobs, W. R., Jr. *Mycobacterium tuberculosis* is Extraordinarily Sensitive to Killing by A Vitamin-C Induced Fenton Reaction. Nat. Commun. 2013, 4, 1881.

(61) Gupta, K.R.; Kasetty, S.; Chatterji, D. Novel Functions of (p)ppGpp and Cyclic di-GMP in Mycobacterial Physiology Revealed by Phenotype Microarray Analysis of Wild-Type and Isogenic Strains of *Mycobacterium smegmatis*. Appl. Environ. Microbiol. 2015, 81, 2571–2578.

(62) Gupta, K.R.; Baloni, P.; Indi, S.S.; Chatterji, D. Regulation of Growth, Cell Shape, Cell Division, and Gene Expression by Second Messengers (p)ppGpp and Cyclic di-GMP in *Mycobacterium smegmatis*. J. Bacteriol. 2016, 198, 1414–1422.

(63) Petchiappan, A.; Naik, S.Y.; Chatterji, D. RelZ-mediated Stress Response in *Mycobacterium smegmatis:* pGpp Synthesis and Its Regulation. J. Bacteriol. 2020, 202, e00444–19.

(64) Boutte, C. C.; Crosson, S. Bacterial Lifestyle Shapes Stringent Response Activation. Trends Microbiol. 2013, 21, 174–180.

(65) Syal, K.; Flentie, K.; Bhardwaj, N.; Maiti, K.; Jayaraman, N.; Stallings, C. L.; Chatterji, D. Synthetic (p)ppGpp Analogue is an Inhibitor of Stringent Response in Mycobacteria. Antimicrob. Agents Chemother. 2017, 61, e00443–17.

(66) Windels, E.M.; Michiels, J.E.; Fauvart, M.; Wenseleers, T.; Van den Bergh, B.; Michiels, J. Bacterial Persistence Promotes the Evolution of Antibiotic Resistance by Increasing Survival and Mutation Rates. ISMEJ. 2019, 13(5), 1239–1251.

(67) Cohen, N.R.; Lobritz, M.A.; Collins, J. J. Microbial Persistence and the Road to Drug Resistance. Cell Host & Microbe 2013, 13(6), 632–642, ISSN 1931-3128.

