## Supplementary data for "*C*-4-Modified Isotetrones Prevent Biofilm Growth and Persister Cell Resuscitation in *Mycobacterium smegmatis*"

### Supporting Information

**Contents** **Page**

Docking and molecular dynamics simulation protocol S-2

NMR spectra S-4

Kill kinetics and persister regrowth S-10

### METHODS

#### Chemical synthesis

Chemicals were purchased from commercial sources. Silica gel (100-200 mesh) was used for purification and TLC analysis was performed on commercial plates coated with silica gel 60 F254 and visualization of the spots was conducted using UV lamp or by spraying 10% polymolybdic acid solution in ethanol. High-resolution mass spectra were obtained from Q-TOF instrument by electrospray ionization (ESI). $^{1}H$ and $^{13}C$ NMR spectral analyses were performed on a spectrometer operating at 400 MHz and 100 MHz, respectively.

#### Docking and molecular dynamics simulation protocol

Rel protein structure was taken from the protein structure database (PDB ID:5XNX), and the missing residues were added using the Modeller 10.3 tool.[1] The structures of ppGpp, Tyrosine Lactone (**12**), and Vitamin C were generated using Avogadro 1.2.0 modelling tool.[2] The blind docking study was performed for all the above mentioned ligands using AutoDock Vina 1.2.3.[3] AutoDock scoring function has contribution from VdW and electrostatic interaction together with H-bonding between protein and ligand.[4] The blind docking was carried out using a grid size of 126×126×126 along the X, Y, and Z axis with 0.708 Å spacing.

AutoDock Vina generated docking conformation with the lowest binding energy was taken as the initial structure for the molecular dynamics (MD) simulation study for each of the protein-ligand complexes. Each protein-ligand complex was solvated using CHARMM modified TIP3P water resulting in a box dimension of 11.02 × 9.98 × 8.80 $nm^3$. The solvated Rel-ppGpp system contained 29736 water molecules, whereas Rel-compound **12** system and Rel-vitamin C system contained 29752 and 29747 water molecules, respectively. Counterions were added to neutralize the systems. Periodic boundary condition was applied in all three directions. All the three protein-ligand systems (Rel-ppGpp, Rel-tyrosine lactone (**12**), Rel-vitamin C) were energy minimized for 16000 xxx steps using the steepest descent algorithm to remove any close contacts. The energy minimized systems are then at first equilibrated at a temperature of 300 K for 1ns in NVT ensemble and then at a pressure of 1bar for 2 ns in NPT ensemble. Modified Berendsen thermostat was used for temperature coupling with a coupling constant of 0.1 ps.[5] Berendsen barostat with a coupling constant of 0.5 ps was used for pressure coupling.[6] Position restraints were applied on the protein-ligand complex during both the equilibrium phases. The equilibrated systems were then subjected to a 10 ns production run, without any position restraints and the coordinate trajectory was recorded for every 10 ps for binding energy calculation. During the production run, Parrinello−Rahman barostat with a coupling constant of 1.0 ps was used for pressure coupling.[7] Integration time was 2 fs during both equilibrium and production run. All bonds containing hydrogen atoms were restrained using LINCS algorithm throughout the simulation time.[8] A cut-off distance of 1.0 nm was used for the computation of the short-ranged L-J interactions and short-range part of the coulomb interactions. Particle mesh Ewald (PME) summation technique was used to calculate long-range coulombic interaction.[9] CHARMM36 all-atom force field with CHARMM- modified TIP3P water model was used to describe the interatomic interactions.[10-12] All simulations were performed in GROMACS 2022.2 package.[13] Binding energy was calculated by means of MM-GBSA method, included in gmx_MMPBSA tool.[14-16] Visualization was carried out using VMD-1.9.3 package, and the calculation of the number of hydrogen bonds was performed using GROMACS module.[17]

**NMR Spectra:**

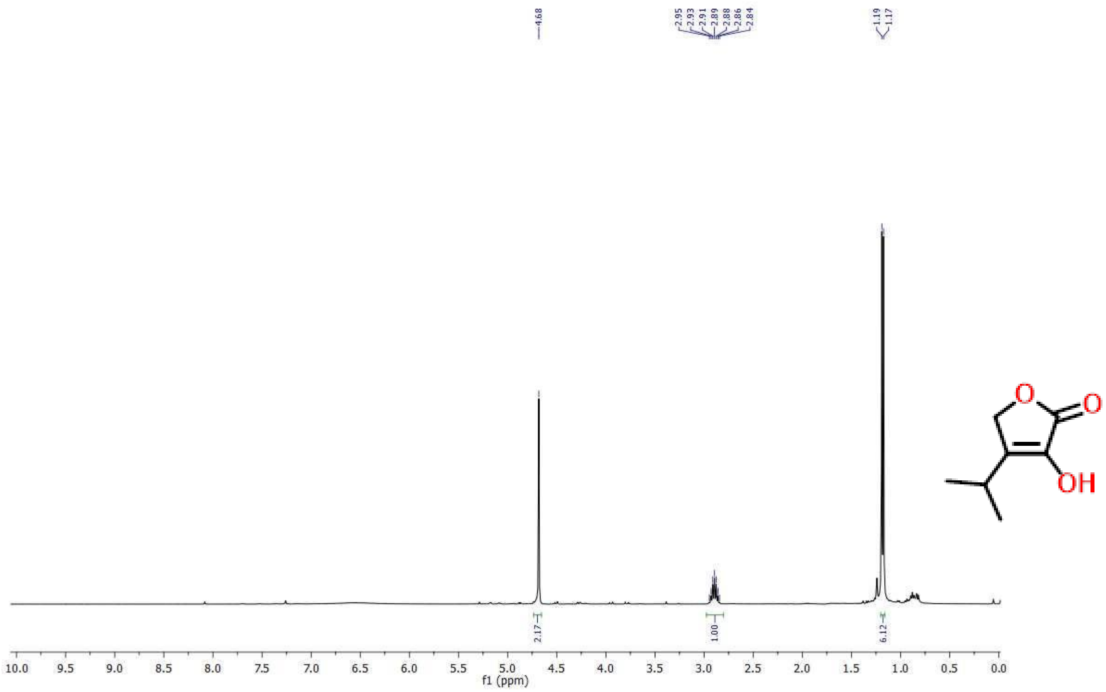


**Figure S1**. $^{1}$H NMR spectrum of **10** ($CDCl_3$, 400 MHz).

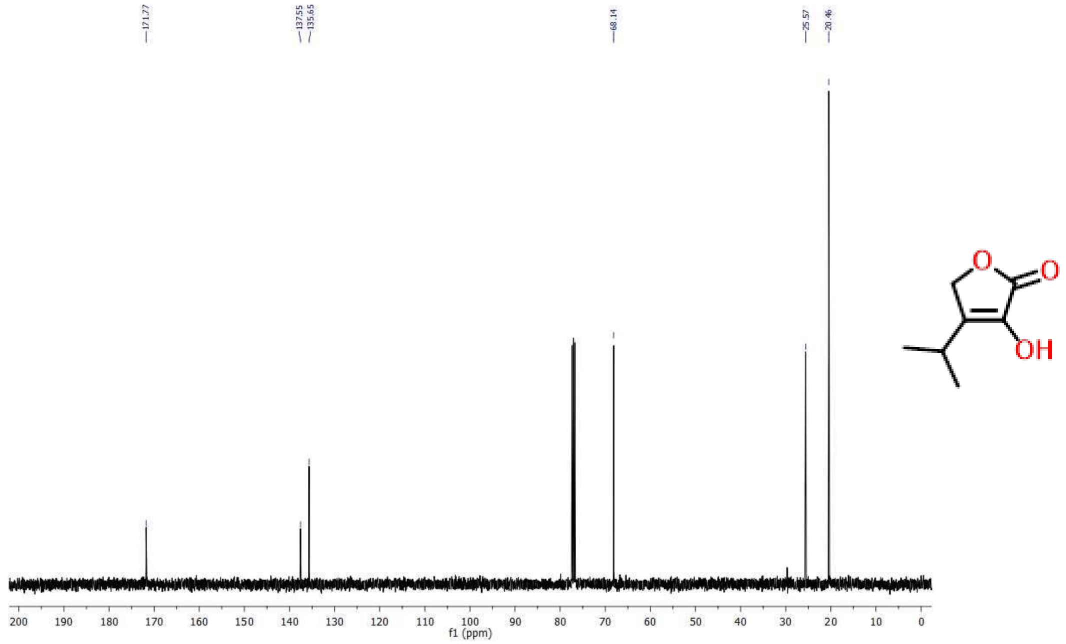


**Figure S2**. $^{13}$C NMR spectrum of **10** ($CDCl_3$, 100 MHz).

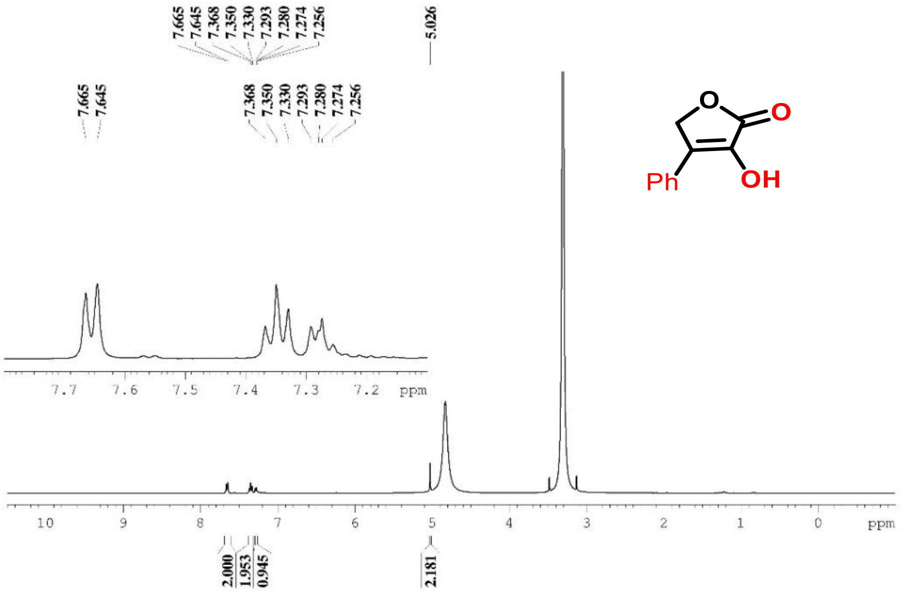


**Figure S3**. [1]H NMR spectrum of **11** ($CD_3OD$, 100 MHz).

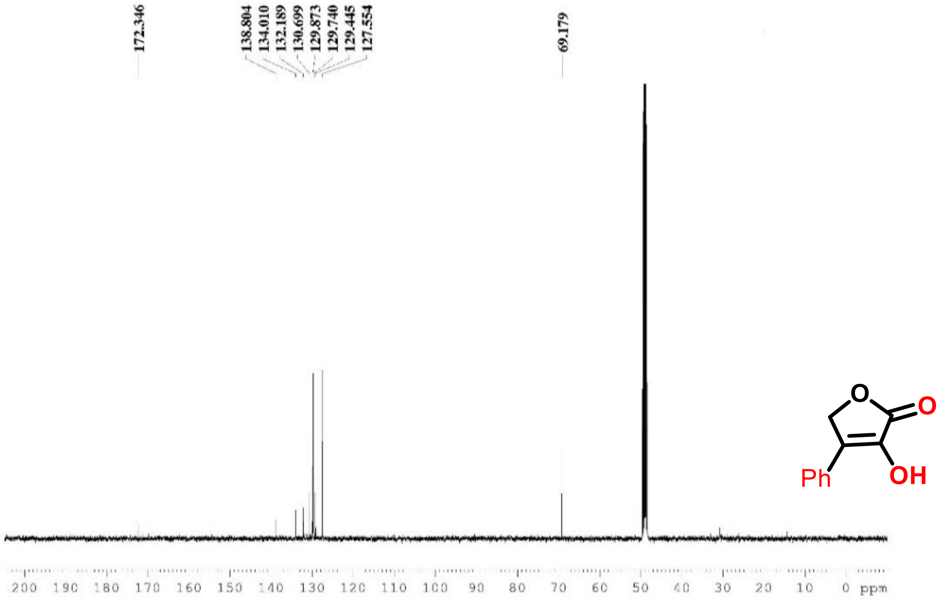


**Figure S4**. [13]C NMR spectrum of **11** ($CD_3OD$, 100 MHz).

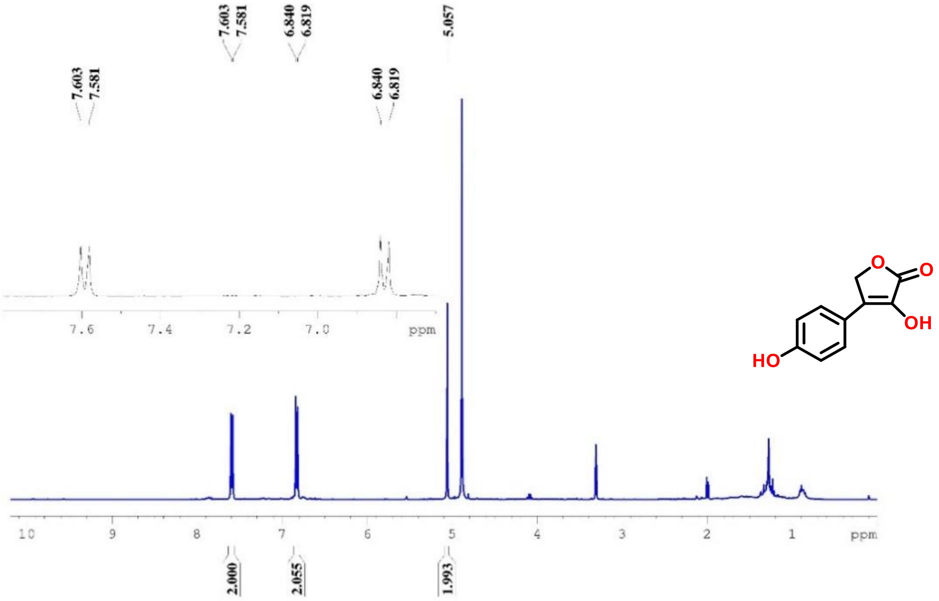


**Figure S5**. $^{1}$H NMR spectrum of **12** ($CD_3OD$, 400 MHz)

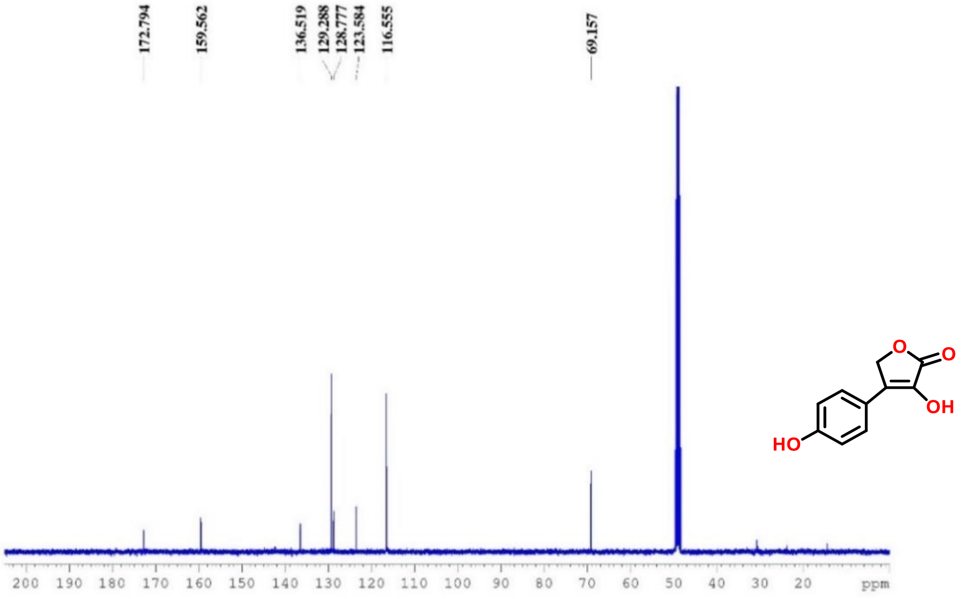


**Figure S6**. $^{13}$C NMR spectrum of **12** ($CD_3OD$, 100 MHz)

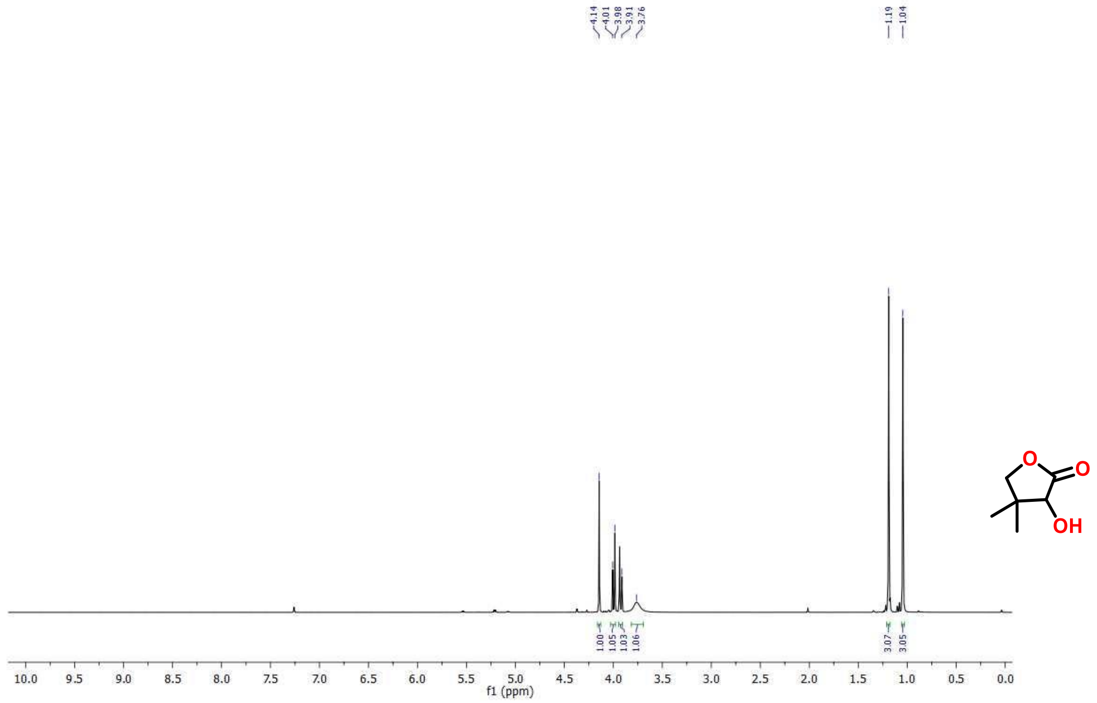


**Figure S7**. [1]H NMR spectrum of **13** ($CDCl_3$, 400 MHz).

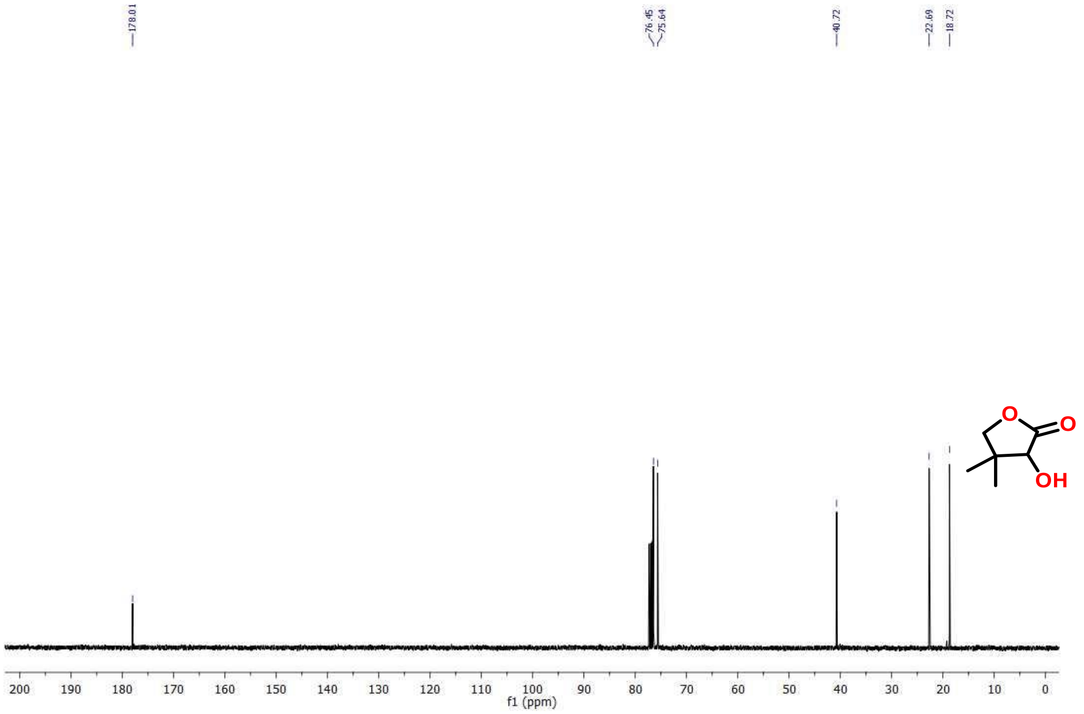


**Figure S8**. [13]C NMR spectrum of **13** ($CDCl_3$, 100 MHz).

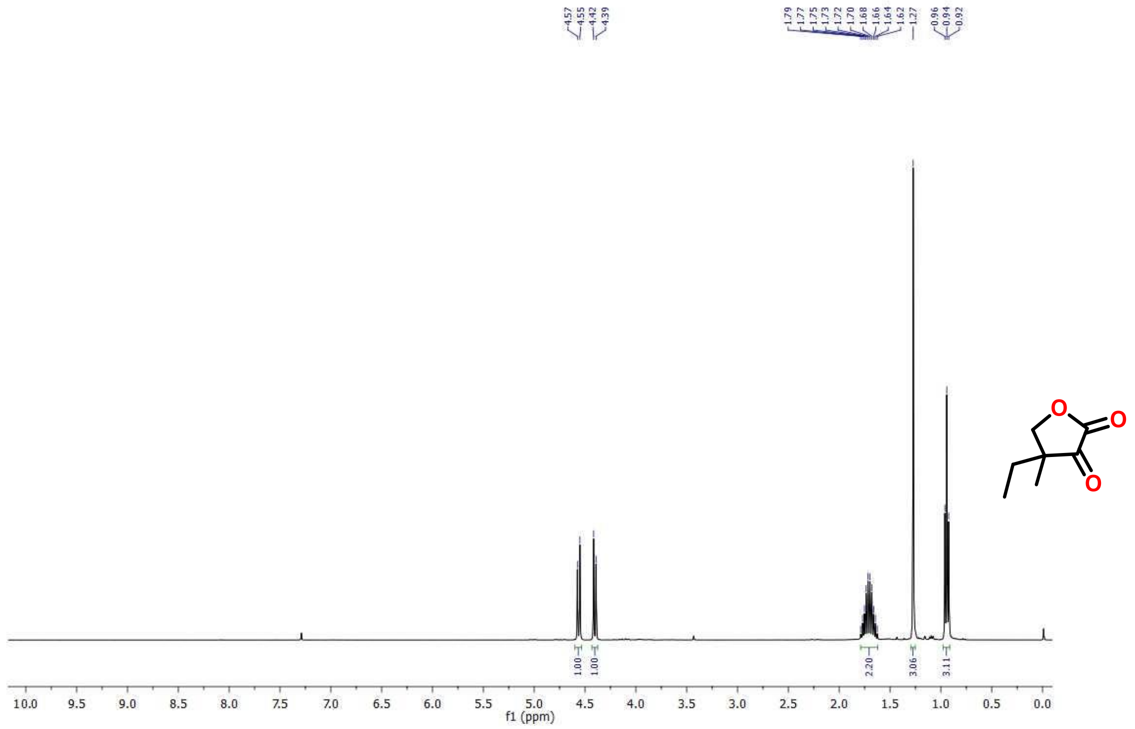

**Figure S9**. $^{1}H$ NMR spectrum of **14** ($CDCl_3$, 400 MHz).

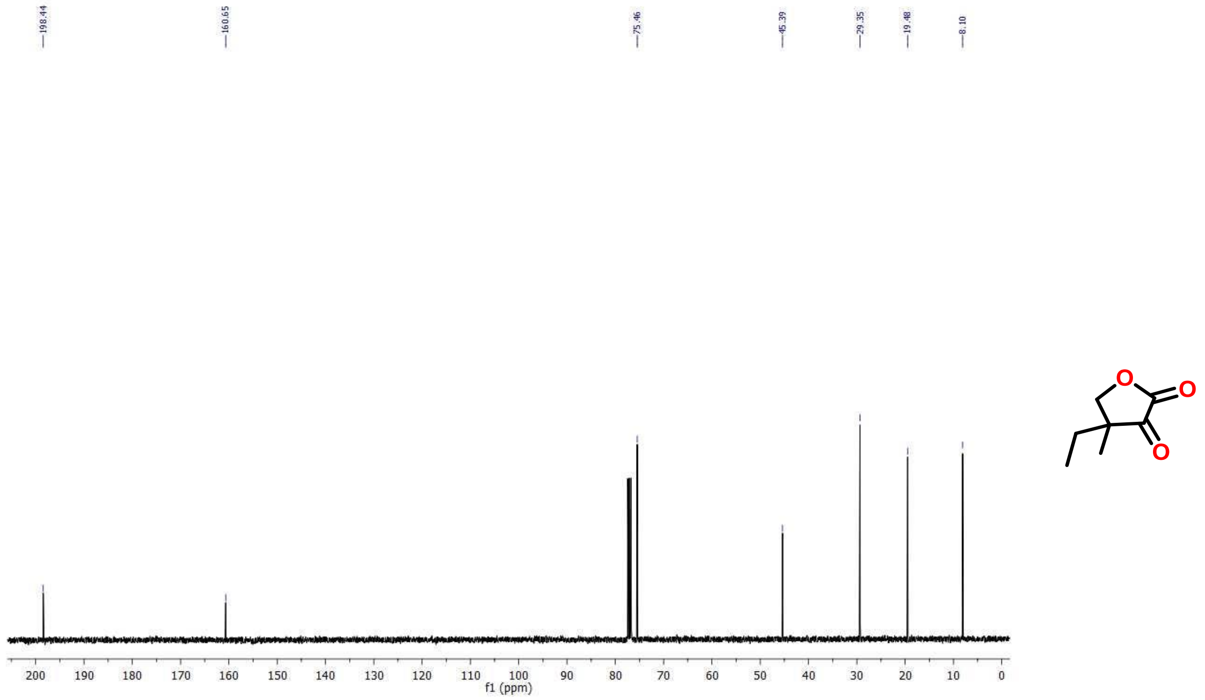

**Figure S10**. $^{13}C$ NMR spectrum of **14** ($CDCl_3$, 100 MHz).

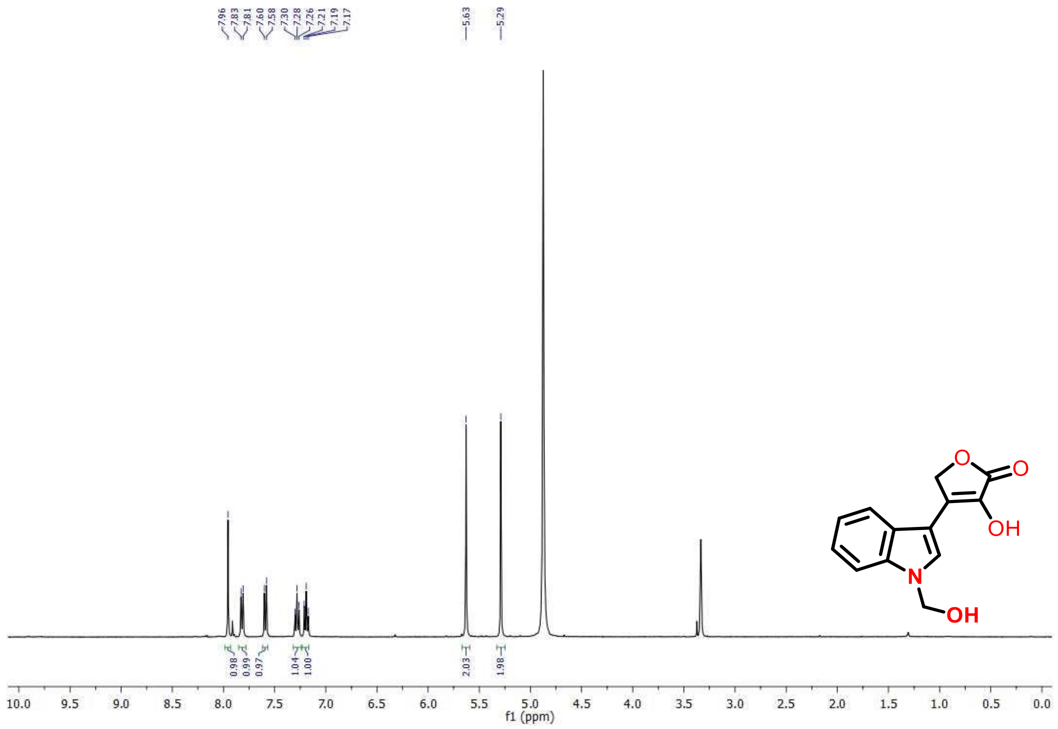


**Figure S11**. $^{1}$H NMR spectrum of **15** ($CD_3OD$, 400 MHz).

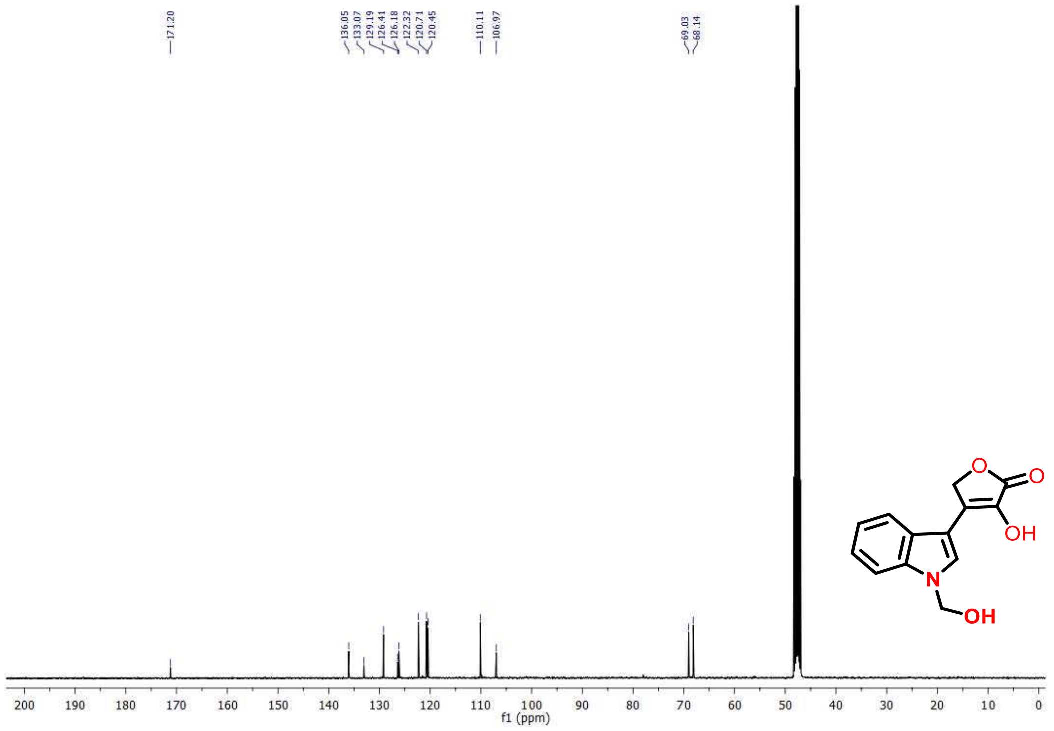


**Figure S12**. $^{13}$C NMR spectrum of **15** ($CD_3OD$, 100 MHz).

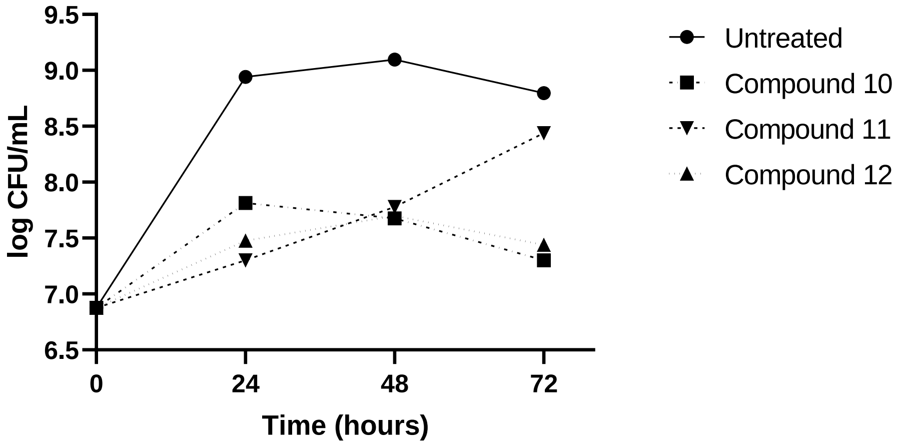


**Figure S13**. Kill kinetics with different isotetrones (without any drug Compounds were used at 400 μg/mL concentration.

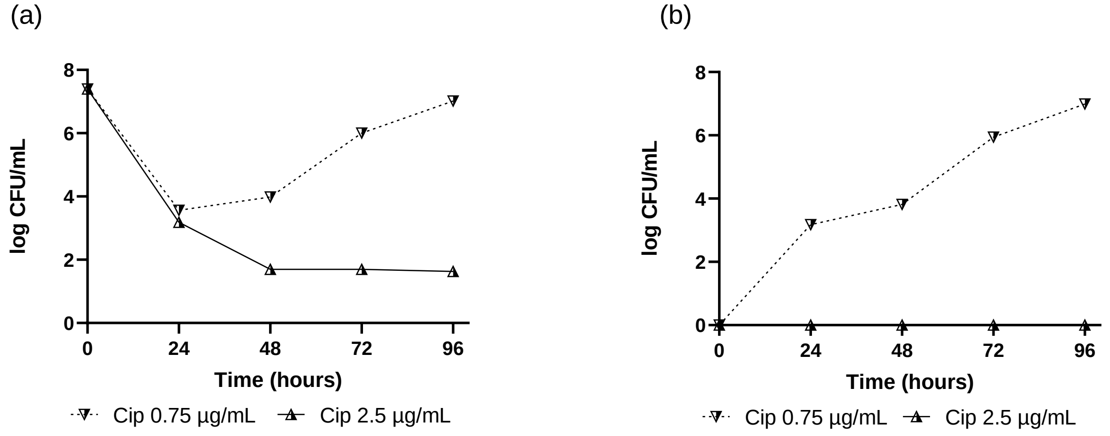


**Figure S14**. Drug tolerant cells' regrowth (a) and resistant mutant enrichment (b) in higher (10X MIC) and lower (3X MIC) concentrations of ciprofloxacin.

### References


(1) Šali, A.; Blundell, T. L. Comparative Protein Modelling by Satisfaction of Spatial Restraints. *J. Mol. Biol.* **1993**, *234* (3), 779–815.

(2) Hanwell, M. D.; Curtis, D. E.; Lonie, D. C.; Vandermeersch, T.; Zurek, E.; Hutchison, G. R. Avogadro: An Advanced Semantic Chemical Editor, Visualization, and Analysis Platform. *J. Cheminform.* **2012**, *4* (1), 1–17.

(3) Trott, O.; Olson, A. J. AutoDock Vina: Improving the Speed and Accuracy of Docking with a New Scoring Function, Efficient Optimization, and Multithreading. *J. Comput. Chem.* **2010**, *31* (2), 455–461.

(4) Morris, G. M.; Huey, R.; Lindstrom, W.; Sanner, M. F.; Belew, R. K.; Goodsell, D. S.; Olson, A. J. AutoDock4 and AutoDockTools4: Automated Docking with Selective Receptor Flexibility. *J. Comput. Chem.* **2009**, *30* (16), 2785–2791.

(5) Bussi, G.; Donadio, D.; Parrinello, M. Canonical Sampling through Velocity Rescaling. *J. Chem. Phys.* **2007**, *126* (1), 14101.

(6) Berendsen, H. J. C.; Postma, J. P. M. van; van Gunsteren, W. F.; DiNola, A.; Haak, J. R. Molecular Dynamics with Coupling to an External Bath. *J. Chem. Phys.* **1984**, *81* (8), 3684–3690.

(7) Parrinello, M.; Rahman, A. Polymorphic Transitions in Single Crystals: A New Molecular Dynamics Method. *J. Appl. Phys.* **1981**, *52* (12), 7182–7190.

(8) Hess, B.; Bekker, H.; Berendsen, H. J. C.; Fraaije, J. G. E. M. LINCS: A Linear Constraint Solver for Molecular Simulations. *J. Comput. Chem.* **1997**, *18* (12), 1463–1472.

(9) Darden, T.; York, D.; Pedersen, L. Particle Mesh Ewald: An N· Log (N) Method for Ewald Sums in Large Systems. *J. Chem. Phys.* **1993**, *98* (12), 10089–10092.

(10) Huang, J.; MacKerell Jr, A. D. CHARMM36 All-atom Additive Protein Force Field: Validation Based on Comparison to NMR Data. *J. Comput. Chem.* **2013**, *34* (25), 2135–2145.

(11) Vanommeslaeghe, K.; Hatcher, E.; Acharya, C.; Kundu, S.; Zhong, S.; Shim, J.; Darian, E.; Guvench, O.; Lopes, P.; Vorobyov, I. CHARMM General Force Field: A Force Field for

Drug-like Molecules Compatible with the CHARMM All-atom Additive Biological Force Fields. *J. Comput. Chem.* **2010**, *31* (4), 671–690.

(12) Neria, E.; Fischer, S.; Karplus, M. Simulation of Activation Free Energies in Molecular Systems. *J. Chem. Phys.* **1996**, *105* (5), 1902–1921.

(13) Abraham, M. J.; Murtola, T.; Schulz, R.; Páll, S.; Smith, J. C.; Hess, B.; Lindah, E. Gromacs: High Performance Molecular Simulations through Multi-Level Parallelism from Laptops to Supercomputers. *SoftwareX* **2015**, *1–2*, 19–25.

(14) Ylilauri, M.; Pentikäinen, O. T. MMGBSA as a Tool to Understand the Binding Affinities of Filamin–Peptide Interactions. *J. Chem. Inf. Model.* **2013**, *53* (10), 2626–2633.

(15) Valdés-Tresanco, M. S.; Valdés-Tresanco, M. E.; Valiente, P. A.; Moreno, E. Gmx_MMPBSA: A New Tool to Perform End-State Free Energy Calculations with GROMACS. *J. Chem. Theory Comput.* **2021**, *17* (10), 6281–6291.

(16) R. Miller, B.; Dwight McGee, T.; M. Swails, J.; Homeyer, N.; Gohlke, H.; E. Roitberg, A. MMPBSA.Py: An Efficient Program for End-State Free Energy Calculations. *J. Chem. Theory Comput.* **2012**, *8* (9), 3314–3321.

(17) Humphrey, W.; Dalke, A.; Schulten, K. VMD: Visual Molecular Dynamics. *J. Mol. Graph.* **1996**, *14* (1), 33–38.
